# Evolved *Bmp6* enhancer alleles drive spatial shifts in gene expression during tooth development in sticklebacks

**DOI:** 10.1101/2021.05.14.444181

**Authors:** Mark D. Stepaniak, Tyler A. Square, Craig T. Miller

**Affiliations:** Department of Molecular and Cell Biology, University of California-Berkeley, Berkeley CA, 94720, USA

**Keywords:** enhancer, *cis*-regulation, evolution, transgene, transgenesis, insulator, fish, stickleback, development, tooth

## Abstract

Mutations in enhancers have been shown to often underlie natural variation but the evolved differences between enhancer activity can be difficult to identify *in vivo*. Threespine sticklebacks (*Gasterosteus aculeatus*) are a robust system for studying enhancer evolution due to abundant natural genetic variation, a diversity of evolved phenotypes between ancestral marine and derived freshwater forms, and the tractability of transgenic techniques. Previous work identified a series of polymorphisms within an intronic enhancer of the *Bone morphogenetic protein 6* (*Bmp6*) gene that are associated with evolved tooth gain, a derived increase in freshwater tooth number that arises late in development. Here we use a bicistronic reporter construct containing a genetic insulator and a pair of reciprocal two-color transgenic reporter lines to compare enhancer activity of marine and freshwater alleles of this enhancer. In older fish the two alleles drive partially overlapping expression in both mesenchyme and epithelium of developing teeth, but the freshwater enhancer drives a reduced mesenchymal domain and a larger epithelial domain relative to the marine enhancer. In younger fish these spatial shifts in enhancer activity are less pronounced. Comparing *Bmp6* expression by *in situ* hybridization in developing teeth of marine and freshwater fish reveals similar evolved spatial shifts in gene expression. Together, these data support a model in which the polymorphisms within this enhancer underlie evolved tooth gain by shifting the spatial expression of *Bmp6* during tooth development, and provide a general strategy to identify spatial differences in enhancer activity *in vivo*.

## INTRODUCTION

The process of development is largely orchestrated by developmental regulatory genes whose spatial and temporal patterns of transcription are controlled by enhancers, *cis*-regulatory elements that bind transcription factors and promote transcription of target genes (Furlong & Levine, 2018; Gasperini et al., 2020). Most developmental regulatory genes are pleiotropic, and function repeatedly at different times and in different tissues during development (Sabarís et al., 2019). Thus, mutations in enhancers of developmental regulatory genes are often more tolerated than coding sequence mutations due to having fewer pleiotropic effects, as the impacts of enhancer mutations are more likely to be restricted in time and/or space, compared to the anatomically more widespread impacts of coding mutations (Carroll, 2008). The importance of enhancers in regulating morphological evolution, natural variation, and disease phenotypes in humans is well established (Rebeiz & Tsiantis, 2017; Rickels & Shilatifard, 2018). However, a growing need has emerged for methods and approaches to compare the activity of molecularly divergent enhancer alleles.

*Cis*-regulatory changes have been shown to underlie the evolution of multiple morphological traits in threespine stickleback fish (*Gasterosteus aculeatus*). Threespine sticklebacks live in both marine and freshwater environments in the Northern Hemisphere, repeatedly forming populations in rivers, streams, ponds, and lakes from ancestral marine populations (Bell & Foster, 1994; McKinnon & Rundle, 2002). Following a freshwater colonization event, a suite of traits has been observed to typically evolve such as reduction in armor (Bell & Foster, 1994; Colosimo, 2005; Cresko et al., 2004) and changes in body shape (Albert et al., 2008; Reid & Peichel, 2010; Walker, 1997; Walker & Bell, 2000). Other traits that typically evolve major differences are those associated with feeding morphology, likely an adaptation to different diets of larger prey in freshwater environments relative to marine ancestral environments (Bell & Foster, 1994; Gross & Anderson, 1984; Hagen, 1967; Lavin & McPhail, 1986; Schluter & McPhail, 1992). High resolution genetic mapping studies have implicated *cis*-regulatory changes as underlying several phenotypes that have evolved in freshwater, including the reduction of armor plates (Archambeault et al., 2020; Colosimo, 2005; Indjeian et al., 2016; O’Brown et al., 2015), pelvic spines (Chan et al., 2010), and pigmentation (Miller et al., 2007), and increases in branchial bone length (Erickson, et al., 2016), and pharyngeal tooth number (Cleves et al., 2014; Cleves et al., 2018).

Increases in pharyngeal tooth number have evolved independently in multiple freshwater stickleback populations (Ellis et al., 2015). Comparing lab-reared marine fish and freshwater fish from the benthic (bottom-dwelling) population of Paxton Lake, revealed that a divergence in tooth number occurs late in development (around ∼20 mm standard length, when fish are juveniles and about half of their adult size). This difference in tooth number continues to increase and becomes more significantly different at adult stages (Cleves et al., 2014). Quantitative trait loci (QTL) mapping identified a large effect QTL that underlies this evolved tooth gain. An F2 cross between a low-toothed Japanese marine fish and a high-toothed benthic Paxton Lake freshwater fish identified a QTL peak on chromosome 21 that explained approximately 30% of the variance in tooth number within the cross. The peak contained the candidate gene *Bone morphogenetic protein 6* (*Bmp6*) which is dynamically expressed in developing teeth. *In situ* hybridization revealed *Bmp6* expression early in the overlying inner, but not outer, dental epithelium (IDE and ODE respectively), as well as in underlying dental mesenchyme, followed by a decrease in expression in the epithelium before the tooth finally erupts into a functional tooth (Cleves et al., 2014; Ellis et al., 2016). Allele specific expression experiments identified *cis-*regulatory changes in *Bmp6*. In tooth tissue from F_1_ hybrids of high-toothed Paxton benthic fish and low-toothed marine fish, a 1.4 fold decrease in *Bmp6* expression from the high-tooth freshwater Paxton benthic allele compared to the marine allele was reported (Cleves et al., 2014). Work in mice and fish has demonstrated an essential role for BMPs in developing teeth (Bei et al., 2000; Cleves et al., 2018; Jia et al., 2013; Vainio et al., 1993; Wang et al., 2012), suggesting a possible causative role of *Bmp6* in evolved tooth gain.

Further refinement of the QTL interval identified a haplotype containing 10 single nucleotide polymorphisms (SNPs) within intron 4 of *Bmp6* that vary concordantly with the presence or absence of the tooth QTL (Cleves et al., 2018). These variable positions define a high-tooth associated haplotype and low-tooth associated haplotype from the Paxton benthic freshwater and marine alleles, respectively. Six core SNPs lie within 468 bases upstream of the previously described minimally sufficient *Bmp6* intron 4 tooth enhancer (Fig. S1) (Cleves et al., 2018). We hypothesized that these core QTL-associated SNPs are modifying the spatial and/or temporal activity of the adjacent tooth enhancer.

Comparing expression patterns of two different alleles of an enhancer through reporter constructs in an organismal context presents two major problems: (1) comparisons of enhancer variants integrated in two different organisms are difficult to fully control for developmental time and genetic background differences and (2) aspects of reporter expression may in part reflect genomic integration site rather than actual enhancer activity. A single bicistronic transgenic construct that contains both enhancer/reporter pairings could address the first problem by providing a comparison within the same animal (and thus both enhancers being compared are at the same stage and in the same genotype). Furthermore, a single bicistronic construct simultaneously reduces the number of genomic integration sites to one and thus reduces position effects, partially addressing the second problem. The placement of a genetic insulator between the enhancer-reporter pairings can reduce cross talk of an enhancer with the opposite paired reporter, creating a more accurate expression profile. Genetic insulators have been shown to be effective in zebrafish (Bessa et al., 2009; Shimizu & Shimizu, 2013). A second alternative approach to a single bicistronic transgene is the use of doubly transgenic two-color lines that include both marine and freshwater enhancers paired with different reporters as parts of separate transgenes. This approach addresses the first problem by having both enhancers in the same animal. With this doubly transgenic two-color line approach, enhancers can be tested with reciprocal pairings (i.e. multiple transgenic reporter lines with different enhancers driving different fluorophores), to control for possible position effects. Here we use transgenic reporter assay experiments to test the hypothesis that the marine and freshwater *Bmp6* intron 4 enhancers have different spatial and/or temporal activity in developing fish embryos, larvae, and adults. We tested this hypothesis in two ways: first, by using a bicistronic enhancer transgene to compare activities of two enhancers in the same fish, and second, by comparing doubly transgenic two-color fish in which the marine and freshwater enhancers drive different fluorophores from different genomic integrations. Lastly, we tested whether the spatial shifts in enhancer activity between marine and freshwater enhancers are also observed for endogenous patterns of *Bmp6* expression during tooth development in marine and freshwater fish.

## MATERIALS AND METHODS

### Animal statement

All animal work was approved by UCB animal protocol #AUP-2015-01-7117-2. Fish were reared as previously described (Erickson et al., 2014).

### Insulator containing bicistronic construct

Gibson assembly was used to create bicistronic constructs to determine insulator efficiency in sticklebacks. Two enhancers with distinct expression domains were used: a 1.3kb fragment from intron 4 of *Bmp6* (Cleves et al., 2018) and the stickleback ortholog of the R2 enhancer for *Col2a1a*, first identified in zebrafish and previously shown to drive similar embryonic expression in sticklebacks (Dale & Topczewski, 2011; Erickson et al., 2016). These two enhancers were placed on opposite sides of a genetic insulator, each with a different reporter gene, either mCherry (mCh) or enhanced GFP (eGFP). The mouse tyrosinase GAB insulator was amplified off the 2pC_GS plasmid (Bessa et al., 2009), while the R2 *Col2a1a* enhancer was PCR amplified from a previously used reporter plasmid (Erickson et al., 2016). The intron 4 enhancer of *Bmp6* was PCR amplified from a reporter plasmid containing either the freshwater allele from the benthic Paxton Lake population or the allele from the Little Campbell marine population (Cleves et al., 2018). All enhancers were PCR amplified simultaneously with the *Hsp70l* promoter as a single amplicon. eGFP and mCh were amplified from previously used reporter plasmids (O’Brown et al., 2015). Primers used and assembly steps are listed in the Supplemental Methods. All components were combined using a Gibson assembly reaction (New England Biolabs ref # E2611L) following the manufacturer’s protocol and transformed into XL1 blue competent cells. Transformed cells were grown on ampicillin containing LB plates and colony inserts were sequence verified by colony PCR. Positive colonies were used to start 50 ml cultures, which were grown overnight. Plasmids were then isolated by Qiagen midi-prep (ref #12145), and Sanger sequence verified.

Tol2 transposase mRNA was transcribed using the plasmid pCS2-TP (Kawakami, 2004) that had been linearized with *Not*I. The linear plasmid was used as template for in vitro transcription using the mMessage SP6 kit (#AM1340). The resulting mRNA was purified using Qiagen RNeasy columns (#74104). Transgene plasmids were co-injected with Tol2 mRNA into newly in vitro fertilized one-cell embryos as described (Erickson et al., 2016). Approximately 200ng of plasmid in 1μl was combined with 1μl of 2M KCl, 0.5μl of 0.5% phenol red, and approximately 1μl of 350 ng/μl of Tol2 transposase mRNA, with water added to a final volume of 5μl, yielding a total concentration of ∼40ng/μl of plasmid and 70ng/μl of mRNA. Embryos were generated from Rabbit Slough (Alaska) marine fish, and lines established and maintained by crossing to lab-reared fish from this same population.

### Generation of single color and doubly transgenic two-color reporter lines

The previously described ∼1.3kb *Bmp6* intron 4 tooth enhancer (Cleves et al. 2018) was amplified from a Paxton Lake benthic fish and Little Campbell marine fish (Figure S1) using the primer pairs MDS35/36 (GCCGGCTAGCGAGAGCATCCGTCTTGTGGG/GCCGGGATCCAGAGTCCTGATGGCCT CTCC) to create reporter plasmids containing the positive orientation (i.e. same 5’ to 3’ orientation as in endogenous locus) of the enhancer relative to the reporter gene or MDS27/28 (GCCGGCTAGCAGAGTCCTGATGGCCTCTCC/GCCGGGATCCGAGAGCATCCGTCTTG TGGG) to create reporter plasmids containing the negative orientation [i.e. the opposite 5’ to 3’ orientation as in the endogenous locus, and possibly more similar to the orientation that an enhancer 3’ to the promoter (e.g. an enhancer in intron 4) would be after looping to contact the promoter] of the enhancer. The fragments were then cloned in both possible 5’ to 3’ orientations into a Tol2 reporter construct upstream of the zebrafish *Hsp70l* promoter and either eGFP or mCherry using *BamH*I and *Nhe*I in the previously generated reporter constructs. Fish that were transgenic for both the marine and the freshwater reporter alleles were generated in one of two ways: (1) crossing of stable lines each containing a single transgene (2) injection of one reporter construct into a stable transgenic line of the opposite (i.e. different population and fluorophore) allele.

### Detecting enhancer activity by fluorescent microscopy

Enhancer activity of the transgenic constructs was imaged by fluorescent microscopy. Previous work demonstrated a *cis*-regulatory difference in *Bmp6* expression between marine and freshwater alleles, with the difference arising late in development (Cleves et al., 2014). As both a divergence in tooth number attributed to the QTL and allele specific expression (ASE) differences arise late in development, post-20 mm total length (Cleves et al., 2014; Cleves et al., 2018), reporter positive fish were dissected at standard lengths pre- and post-tooth number divergence (20 mm total length) as previously described (Ellis & Miller, 2016). Tooth plates were then fixed in 4% PFA in 1x PBS for 60 minutes, washed through a graded series of 3:1, 1:1, 1:3 water and glycerol solutions into 100% glycerol, flat-mounted, and imaged. Comparisons were made across the different alleles and orientations on a Leica M165FC fluorescent dissecting microscope with filters GFP1 (#10447447) and RhodB (#10447360), and a Leica DM2500 compound microscope with filters GFP (#11532366) and TX2 (#11513885). To compare enhancer activity in fish before and after tooth divergence (20 mm standard length), ventral tooth plates and dorsal tooth plates were imaged and enhancer activity was assessed in the dental epithelium and mesenchyme of each tooth, in each of three pre-divergence sized fish (between 16 - 18.5 mm total length) and three post-divergence sized fish (between 30 – 48 mm total length) in two different sets of integrations and enhancer/reporter pairings. If the QTL-associated SNPs are responsible for the QTL peak and therefore tooth number differences observed late in development, as well as the ASE differences, we would expect the enhancers to have different activity in > 20 mm fish compared to < 20 mm fish. We would also expect the enhancers to have similar activity earlier in development, when allele specific expression was not significantly different between the freshwater and marine alleles (Cleves et al., 2014).

### Quantification of enhancer activity differences across tooth development

As we hypothesized that the QTL associated intronic polymorphisms result in differential enhancer activity in the dental mesenchyme and/or epithelium, we characterized enhancer activity in both tissues across multiple tooth plates. The stage of each tooth was scored as either early (late cap to early bell stages in which mesenchyme has condensed under the epithelium but no mineralization has occurred), middle (mineralization of the forming tooth has started to occur, also called late bell stage) or late (a fully formed tooth has erupted, also called functional stage (Ellis et al., 2015)). The activity for each enhancer allele was recorded as either present or absent in the epithelium (early and middle stages) and mesenchyme. Additionally, we also recorded if either allele (marine or freshwater) drove more robust or extensive expression in each domain, indicating an allelic bias.

### *In situ* hybridization on sections

Stickleback adult (∼40 cm standard length) pharyngeal tissues were prepared, sectioned, and assayed by ISH in parallel to compare the spatial distribution of *Bmp6* mRNA. Adults derived from marine (Rabbit Slough [RABS]) and freshwater (Paxton Benthic [PAXB]) populations were euthanized, and their pharyngeal tissues were fixed overnight in 4% formaldehyde (Sigma P6148) in 1x phosphate-buffered saline (PBS) at 4° C with heavy agitation, washed 3x 20 min with PBST on a nutator, then decalcified for 5 days in 20% ethylenediaminetetraacetic acid (EDTA, pH 8.0) at room temperature on a nutator. Marine and freshwater fish were always collected and prepared in parallel such that all storage and preparation intervals were equivalent. The *in situ* hybridization (ISH) for *Bmp6* was carried out as described previously (Square et al., 2021), with some modifications to ensure maximally comparable assays were carried out on marine and freshwater samples in parallel. A previously published *Bmp6* riboprobe was used in this study (Cleves et al., 2014; Square et al., 2021). The *Bmp6* riboprobe was synthesized with digoxygenin-labeled UTP and added at a concentration of ∼300 ng/mL in 20 mL of hybridization buffer, split between 2 different LockMailer slide containers (Sigma-Aldrich), and agitated overnight in a rotating hybridization oven at 67° C. Slides from marine and freshwater fish were cohoused in the hybridization buffers to ensure equal exposure to the riboprobe between marine and freshwater samples. Hybridization buffer washes, blocking, and antibody incubation steps were as previously described (Square et al., 2021). Signal development was carried out for 2, 3, or 7 days to visualize mRNA localization. Marine and freshwater slides were developed in parallel (in the same solutions, in the same LockMailer containers), and only those sections that experienced the same coloration reaction were compared (i.e. we only directly compared sections that were prepared in parallel). To prepare slides for imaging, they were counterstained with DAPI, rinsed then washed 3x 5+ min with deionized H_2_0, coverslipped with deionized H_2_0, and imaged on a Leica DM2500 microscope. The procedure outlined in this section was replicated three times, each replication used two marine and two freshwater adults, for a total of n=6 fish from each background.

## RESULTS

### Two ways to compare enhancers in transgenic fish

We used two strategies to compare enhancer alleles in the same transgenic fish. First, we used a single bicistronic construct with a genetic insulator separating two enhancer/reporter pairs. Second, we used two separate transgenic constructs, independently integrated in the same fish line and each containing a single enhancer allele (marine or freshwater, Figure S1) with a distinct fluorescent reporter (eGFP or mCherry), to generate doubly transgenic two-color fish.

### Insulator efficiency in F_0_ fish

To test the first strategy of a bicistronic construct separated by an insulator, a bicistronic construct was generated using two enhancers that drive expression in non-overlapping domains. In sticklebacks, the *Col2a1a R2* enhancer drives expression in the developing notochord with expression seen by the third day post fertilization (dpf) (Erickson, Ellis, et al., 2016). By 8 dpf we observed *R2* reporter expression in the developing craniofacial skeleton, including Meckel’s cartilage, the hyosympletic, and the ceratohyal (Figure S2), similar to the reported enhancer activity in zebrafish (Dale & Topczewski, 2011). The *Bmp6* intron 4 tooth enhancer has not been reported to drive expression in the domains seen in the *R2 Col2a1a* enhancer. In addition, the previously described tooth and early fin domains (Cleves et al., 2018), as well as the presently described late fin domains, are not domains in which the *Col2a1a* enhancer has been observed to drive expression. Thus, to our knowledge these two enhancers drive distinct and non-overlapping expression domains within these embryonic and larval tissues, providing multiple locations that can test for insulation within the construct.

Three clutches were injected with a *Col2a1a* enhancer/*Bmp6* tooth enhancer bicistronic construct (Figure 1A) for a total of 228 injected embryos, of which 92 were scoreable at 7 dpf. Four domains (left and right pectoral fins, median fin fold, and notochord) were scored for insulation efficiency (0-2 for no to complete insulation, see Supplemental Methods). Across all domains the average insulator score was 0.94 (Table S1). Overall, the bicistronic construct using the mouse tyrosinase insulator element (GAB) moderately prevented reporter genes from being activated by nearby enhancers when placed between the elements. Within the same F_0_ fish we observed both insulated and uninsulated domains, with insulation even varying within a domain (Figure 1B). For example, insulation was observed in the median fin and left pectoral fin, but not within some regions of the right pectoral fin of a 7 dpf embryo in which both mCherry and eGFP were observed. To control for enhancer/reporter pairing, the inverse construct was created, with the *Col2a1a* enhancer driving eGFP and the *Bmp6* tooth enhancer driving mCherry. A total of 154 fish were injected across two clutches, with 30 surviving to 7 dpf that were scoreable, with an average score of 0.64 (Table S2). Overall, both insulator constructs demonstrate the ability to drive some degree of separate expression domains of two enhancers concurrently, consistent with results reported in zebrafish that showed insulators can block enhancer-promoter crosstalk (Bessa et al., 2009).

**Figure 1.**
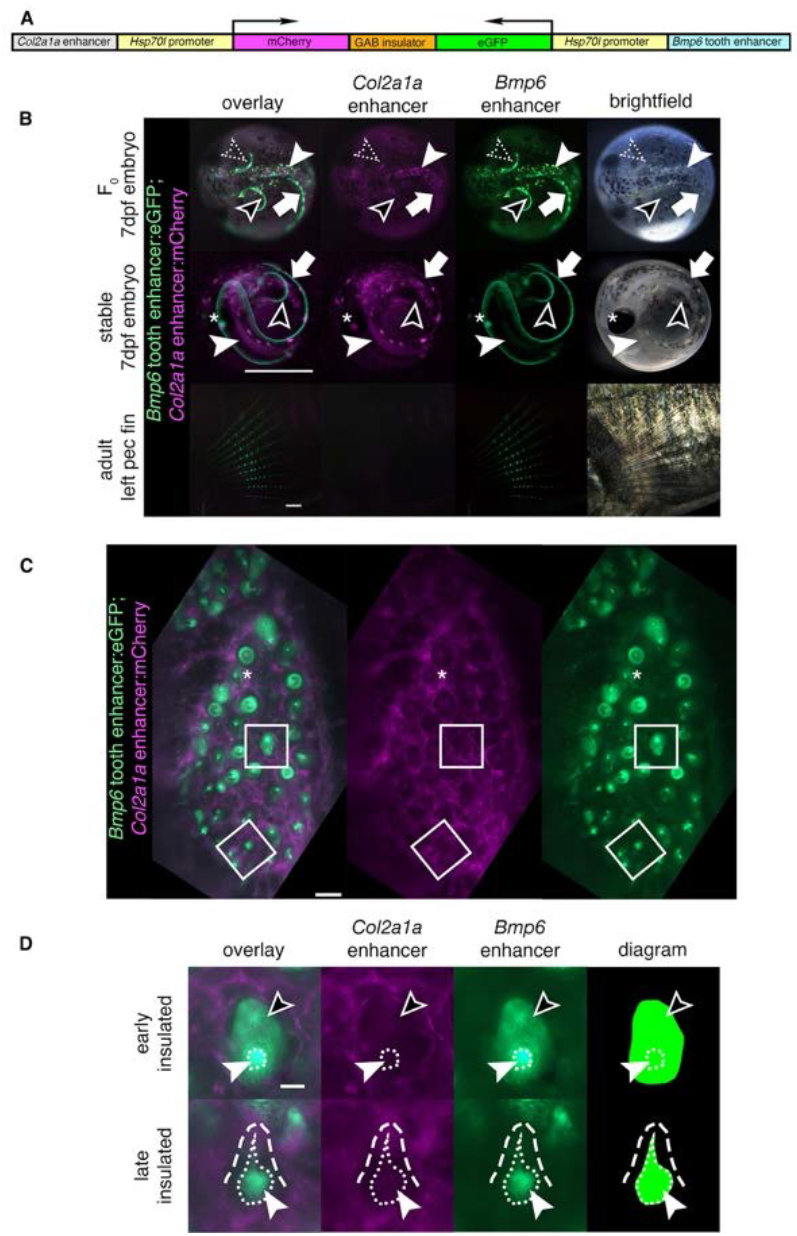
An insulated bicistronic construct reports separate expression patterns from two different enhancers. (**A**) Bicistronic construct with a *Col2a1a* enhancer and *Hsp70l* promoter driving mCherry and the freshwater *Bmp6* intronic tooth enhancer and *Hsp70l* promoter driving eGFP, separated by the mouse tyrosinase insulator (GAB). (**B**) Transgenic fish show a separation of domains in red and green overlay, red channel only, green channel only, and brightfield (left to right). Top: In 7 days post fertilization (dpf) F_0_ fish (dorsal view), insulation was observed in some but not all domains. Both mCherry and eGFP were observed in the same area in the right pectoral fin (dotted arrowhead), indicating incomplete or failed separation of domains, while in the other areas of the pectoral fin only eGFP was observed (black arrowhead). Within the notochord (solid white arrowhead), only mCherry was observed, while in the median fin (white arrow) only eGFP was observed, indicating insulation in both domains. Middle: In 7 dpf stable F_1_ fish (lateral view), only eGFP was observed in the pectoral fins (black arrowhead) indicating successful insulation in those domains, while both fluorophores were detected in the median fin (white arrow) and in the notochord (solid white arrowhead) indicating a lack of insulation. Both fluorophores were detected in the lens of the eye (asterisk), a domain driven by the *Hsp70l* promoter. Bottom: in adult pectoral fins (lateral view), eGFP but not mCherry expression was detected. (**C-D**) Dorsal pharyngeal tooth plate (C) and representative teeth of early and late stages (D) from adult stable transgenic fish. (C) Insulator effectiveness was observed with eGFP restricted to predicted tooth domains and mCherry primarily present in the surrounding tissue. In some teeth, faint mCherry appeared to be expressed in the dental mesenchyme (asterisk). (**D**) eGFP expression was detected in the dental mesenchyme (solid arrowhead and extent of mesenchyme as white dotted line) and dental epithelium (black arrowhead) of developing teeth, while mCherry was expressed in the surrounding tissue (white dashed line outlines a mineralized tooth). Scale bars = 1mm (**B)**, 100μm (**C),** 25μm (**D**).

### Insulator effectiveness in stable fish

Variation in insulator effectiveness across an individual F_0_ fish may be due to different genomic integrations of the bicistronic constructs. To determine the effectiveness of a single bicistronic transgene, F_0_ fish were outcrossed to create stable F_1_ individuals for the *Col2a1a R2*:mCherry; *Bmp6* tooth enhancer:eGFP bicistronic construct. In 7 dpf F_1_ embryos, complete fin domains of the *Bmp6* enhancer were observed, with insulation apparent in some but not all domains (Figure 1B). In adults, *Bmp6* enhancer activity was observed in the intersegmental joints of fins (described below), however no mCherry was observed, suggesting effective insulation in that domain (Figure 1B). Insulator activity was also observed in pharyngeal teeth (Figure 1C). The *Bmp6* enhancer was observed to drive expression in the mesenchyme and inner dental epithelium (IDE) of pharyngeal teeth (Figure 1D), consistent with previous reports. mCherry was not observed in the tooth domains, suggesting effective insulation in adult teeth. Thus, in stable transgenic adults the insulator can separate the activity of the two enhancers, including within the dental epithelium and mesenchyme domains of the *Bmp6* enhancer.

### Bicistronic construct reveals spatial shifts in mesenchymal and epithelial activity of *Bmp6* enhancer alleles

Since the GAB genetic insulator can block enhancer-promoter crosstalk in bicistronic constructs, a bicistronic construct with both the marine and freshwater alleles (Figure 2A) was used to create a stable line as a first test for enhancer activity differences. The marine allele, paired with mCherry, appeared to drive a more robust mesenchymal domain compared to the freshwater allele (Figure 2B-C). In contrast, within the inner dental epithelium more GFP than mCherry signal was detected, suggesting an expanded epithelial domain driven by the freshwater enhancer compared to the marine allele. Thus, in developing teeth from fish with this bicistronic transgene, the marine allele drove more robust expression in the mesenchyme while the freshwater allele drove more robust expression in the epithelium.

**Figure 2.**
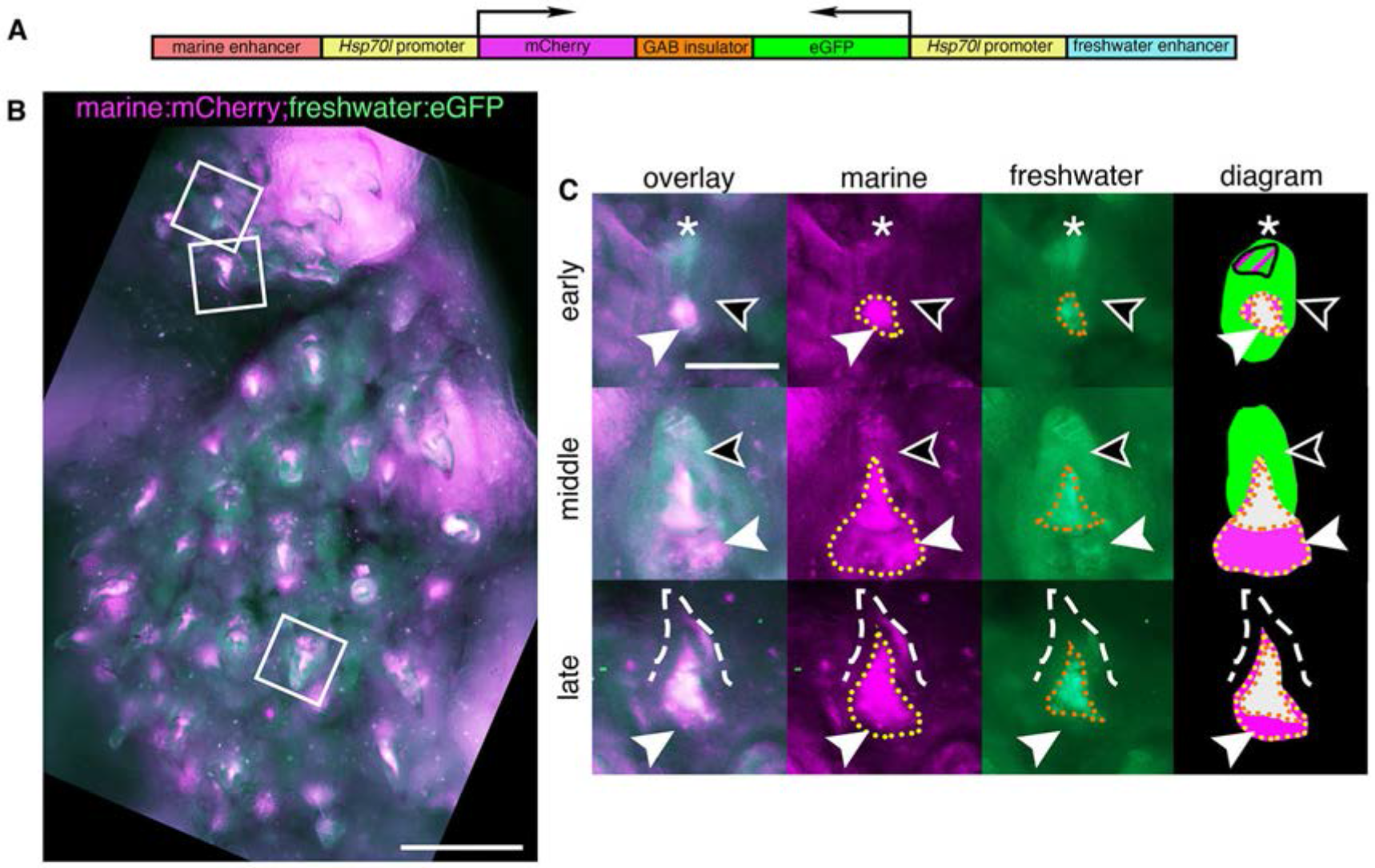
A bicistronic construct using a genetic insulator separates the expression domains of the marine and freshwater alleles of the *Bmp6* tooth enhancer. (**A**) Bicistronic construct with the marine allele of the intron 4 *Bmp6* enhancer/*Hsp70l* promoter driving mCherry and the freshwater allele/*Hsp70l* promoter driving eGFP separated by the mouse tyrosinase GAB insulator. (**B**) Dorsal pharyngeal tooth plate from a fish transgenic with construct (A), and representative teeth (white boxes) from early, middle, and late stages (early bell, late bell, and functional, respectively) (**C**). Early: epithelium expressed eGFP throughout (black arrowhead) while a concentrated tip (asterisk) was observed to contain both marine and freshwater activity. In the mesenchyme (white arrowhead) the marine allele had a more robust and larger expression domain (yellow dotted line) compared to the freshwater allele (orange dotted line). Middle: epithelium had freshwater expression while the marine allele continued to drive more robust expression in the mesenchyme compared to the freshwater allele. Late: As in the other stages the freshwater allele had a more restricted expression domain in mesenchyme of erupted mineralized teeth (dashed line). Scale bars = 200μm (**B),** 50μm (**C)**.

### Doubly transgenic fish confirm expanded freshwater epithelial *Bmp6* enhancer activity in post-divergence fish

As a second method to compare the spatial and temporal activity of marine and freshwater enhancer alleles, we generated stable bi-color transgenic lines with the two different alleles of the *Bmp6* intron 4 tooth enhancer on separate constructs: freshwater:eGFP;marine:mCherry, in the opposite 5’ to 3’ direction as the endogenous locus, and freshwater:mCherry;marine:eGFP, in the same 5’ to 3’ direction as the endogenous locus. In adult fish, both marine and freshwater enhancers were observed to drive dynamic expression in the IDE, more intensely at earlier stages, and diminishing as development of the tooth approaches eruption (Figure 3A-C, and Figure S3A-C), consistent with *Bmp6* expression detected by whole-mount in situ hybridization (Cleves et al., 2014; Ellis et al., 2016). In multiple tooth germs, a brighter focus was observed at the distal tip of the epithelium with both enhancers (Figure 3A-C & Figure S3A-C), a domain resembling the localized distal epithelial expression of *Fgf10* and putative enamel knot in shark embryos (Rasch et al., 2016). This distal epithelial domain was the last epithelial region to drive reporter expression prior to cessation in the epithelium. While both enhancers were observed to drive expression in the epithelium, the freshwater allele drove seemingly more robust expression of the reporter, both in terms of intensity as well as spatial extent of the domain (Figure 3B-C, Figure S3B-C).

**Figure 3.**
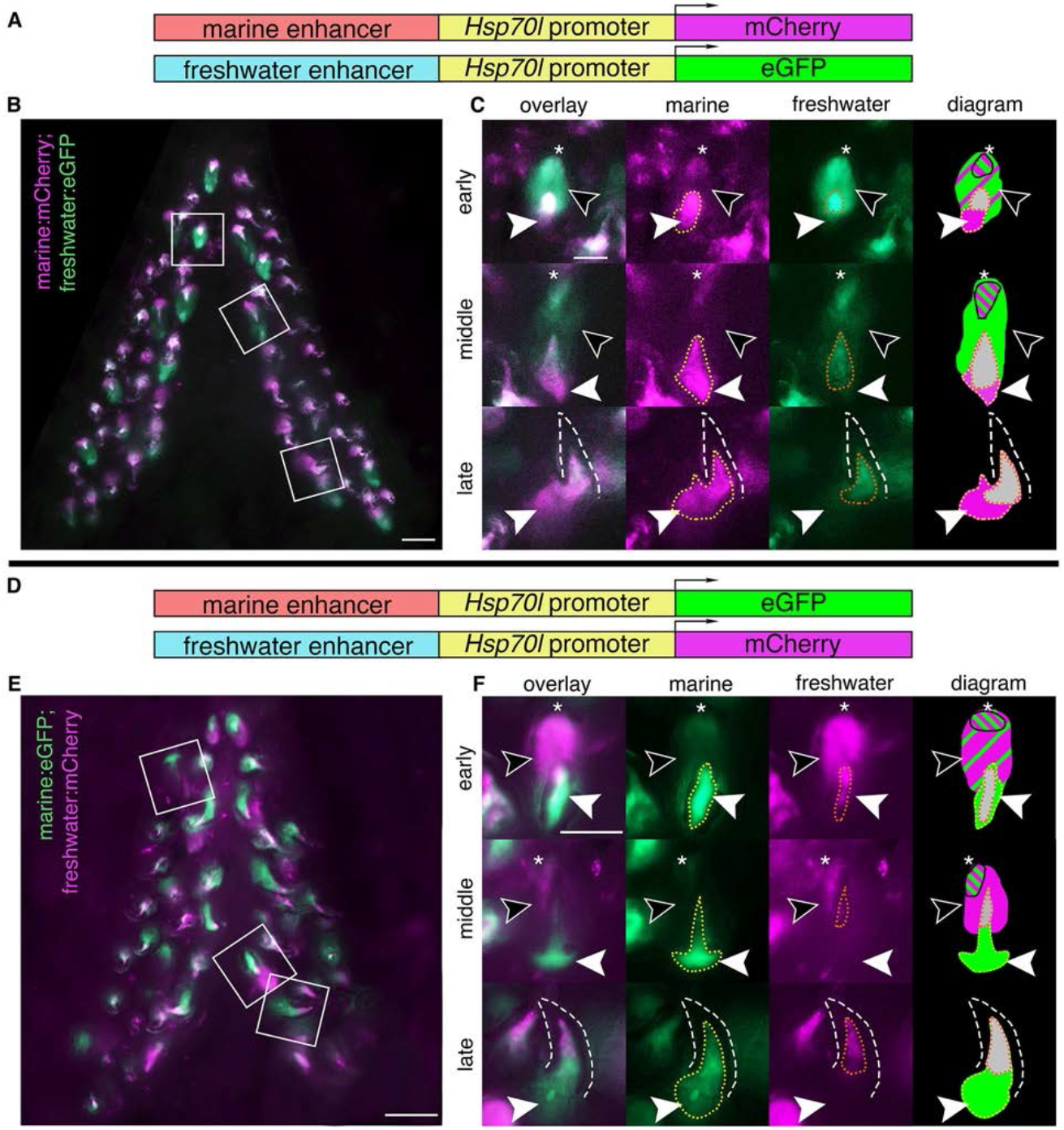
Reduced mesenchymal and expanded epithelial expression of freshwater enhancer relative to marine enhancer in developing teeth. Ventral pharyngeal tooth plates from fish doubly transgenic for two alleles of the *Bmp6* intron 4 enhancer driving two different reporter genes (**A,D**): the marine enhancer driving mCherry with the freshwater enhancer driving eGFP (**B,C**) and the marine enhancer driving eGFP with the freshwater enhancing driving mCherry (**E,F**). Bilateral ventral pharyngeal tooth plates (**B,E**) are shown, next to representative teeth from three stages (**C,F**): early (early bell), middle (late bell), and late (functional) highlighted by white boxes in **B,E**. (**C,F**) Early: freshwater and marine enhancer drove expression in the epithelium (black arrowheads), with concentrated expression in the tip (asterisk), and more overall epithelial expression from the freshwater enhancer. Both enhancers also drove expression in the mesenchyme (solid white arrowhead) with a larger expression domain of the marine allele (yellow dotted line) compared to the freshwater allele (orange dotted line) seen in both genotypes. Middle: freshwater allele still drove expression in the epithelium while marine allele had reduced or undetectable expression outside concentrated tip. The marine allele drove more robust mesenchymal expression compared to the freshwater allele. Late: marine allele drove robust expression in the mesenchyme compared to freshwater allele in mineralized tooth (dashed line). Diagram: summary of tooth epithelial and mesenchymal domains. The relative sizes of green and magenta hatched lines correspond to the approximate relative strength of expression in the epithelium. Overlapping mesenchyme domain is grey, and expanded marine mesenchyme is marked with white arrowhead. Scale bars = 100μm (**B**,**D)**, 50μm (**C**, **F)**.

### Doubly transgenic fish confirm reduced freshwater mesenchymal *Bmp6* enhancer activity in post-divergence fish

Reporter expression from the two alleles appeared in the mesenchyme of teeth across all stages. In pre-eruption (early and middle stage) tooth germs, condensed mesenchyme was observed to show activity of both enhancers (Figure 3B-C and Figure S3B-C). In fully formed, erupted, late-stage teeth, reporter expression was observed in the mesenchymal core, extending from the tip of the core down to the base of the tooth where expression widened. Deeper mesenchyme was observed to consistently display marine but not freshwater enhancer activity. The deeper, broader, and more robust mesenchymal expression domain driven by the marine allele compared to the freshwater allele was also observed in stages of tooth development prior to eruption (Figure 3B-C and Figure S3B-C).

### Reciprocal reporter/enhancer pairing in second doubly transgenic two-color line support epithelial and mesenchymal shifts in enhancer activity

To determine if the previous observations were artifacts due to factors such as transgene position effects, fluorophore used, or enhancer orientation, next we made constructs where each enhancer had an opposite enhancer orientation and drove the other fluorophore (Fig. 3D). These constructs were then randomly integrated by Tol2-mediated transgenesis, representing independent genomic integrations of oppositely oriented enhancers with alternate fluorophores, simultaneously controlling for genomic position effect, enhancer orientation, and fluorophore strength. Using these reciprocal constructs, we again observed the epithelial and mesenchymal differences seen in the bicistronic construct and the first double transgenic line, suggesting the QTL-associated freshwater SNPs reduce mesenchymal and expand epithelial enhancer activity (Figure 3E-F and Figure S3E-F).

### Less pronounced enhancer activity differences in early fish

Allele specific differences in the expression levels of the freshwater and marine alleles of *Bmp6*, as well as tooth number, have been shown to arise later in development (> 20 mm fish length). We hypothesized that if the SNPs found within the freshwater and marine haplotypes contribute to the allele specific expression differences, and subsequently tooth number differences, the differences in enhancer expression should be more pronounced in larger fish compared to smaller fish. Fish smaller than the tooth divergence point (∼16-18.5 mm juveniles, see Methods) were dissected from each genotype and tooth plates were fixed and imaged (Figure 4). While the epithelial and mesenchymal expression differences observed in the older post-divergence stages were still present in both the dental epithelium and mesenchyme (Figure 4C,F), the enhancer differences were less pronounced. In multiple early and middle stage teeth the epithelium showed similar activity from both alleles (Figure 4C,F), unlike the expanded freshwater epithelial domain that was observed in larger fish. Overall, the expression patterns of the two enhancers appeared more similar in pre-divergence fish, consistent with previous allele specific expression and tooth number results (Cleves et al., 2014).

**Figure 4.**
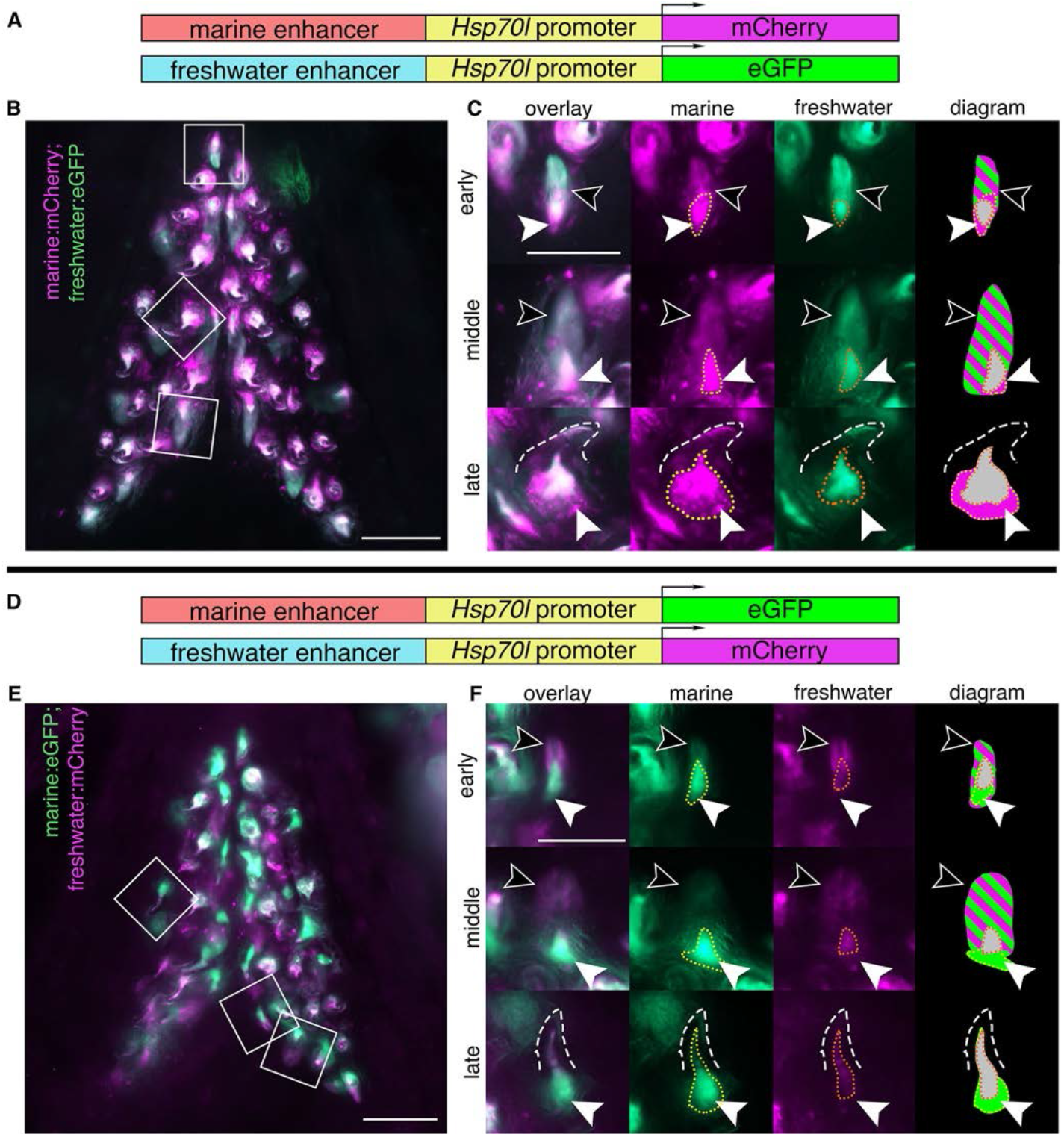
Marine and freshwater *Bmp6* enhancers drive more similar spatial patterns in younger fish. Ventral pharyngeal tooth plates from < 20 mm (pre-tooth number divergence) fish doubly transgenic for two alleles of the *Bmp6* intron 4 enhancer driving two different reporter genes (**A,D**): the marine enhancer driving mCherry with the freshwater enhancer driving eGFP (**B,C**) and the marine enhancer driving eGFP with the freshwater enhancing driving mCherry (**E,F**). Bilateral ventral tooth plates (**B,E**) are shown next to representative teeth from the three stages (**C,F**): early, middle, and late highlighted by white boxes in **B,E**. Early: both freshwater and marine enhancer drove expression robustly in the epithelium (black arrowheads), while both enhancers drove expression in the mesenchyme (white arrowheads), the marine enhancer drove a broader domain (yellow dotted line) compared to the freshwater enhancer (orange dotted line). Middle: both enhancers continued to drive robust, apparently similar levels of expression in the epithelium (black arrows). In the mesenchyme (white arrowheads) the domain of the freshwater enhancer was reduced compared to the marine allele. Late: marine allele continued to drive a broader domain within the mesenchyme of mineralized teeth (dashed line). The relative sizes of green and magenta hatched lines correspond to the approximate relative strength of expression in the epithelium. Overlapping mesenchyme domain is grey, and expanded marine mesenchyme is marked with white arrowhead. Scale bars = 100μm (**B**,**E)**, 50μm (**C**, **F)**.

### Quantification of epithelial and mesenchymal expression patterns

Quantification of epithelial and mesenchymal expression, and bias towards enhancer activity was scored for three tooth plates of each type (ventral and dorsal) at pre and post tooth number divergence (Supplemental Material). In post divergence fish, activity of the freshwater enhancer was observed in the epithelium in both ventral and dorsal tooth plates in nearly all pre-eruption teeth (Figure 5A & Table S3). The marine allele was detected in the epithelium of only a subset of pre-eruption teeth, from approximately 70-90% of pre-eruption teeth in pooled tooth plate data (Figure 5A). When combining tooth plate data for each genotype the marine enhancer was active in the epithelium in a higher percentage of early stage germs compared to middle stage (marine:mCherry;freshwater:eGFP early: 44/52 [84.6%], middle 39/51 [76.5%] and marine:eGFP;freshwater:mCherry early: 39/47 [83.0%], middle 30/40 [75%]). The pattern is still present when data is sorted by tooth plate and genotype (Supplemental Material). Therefore, while there does appear to be a stage effect, variation also exists within stages. Overall, the freshwater enhancer drove expression more frequently and more robustly in the epithelium of early and middle stage teeth compared to the marine allele in post divergence fish. However, in pre-divergence fish, the epithelium of all pre-eruption teeth exhibited robust expression of both enhancers, across both genotypes and tooth plates (Figure 5A).

**Figure 5.**
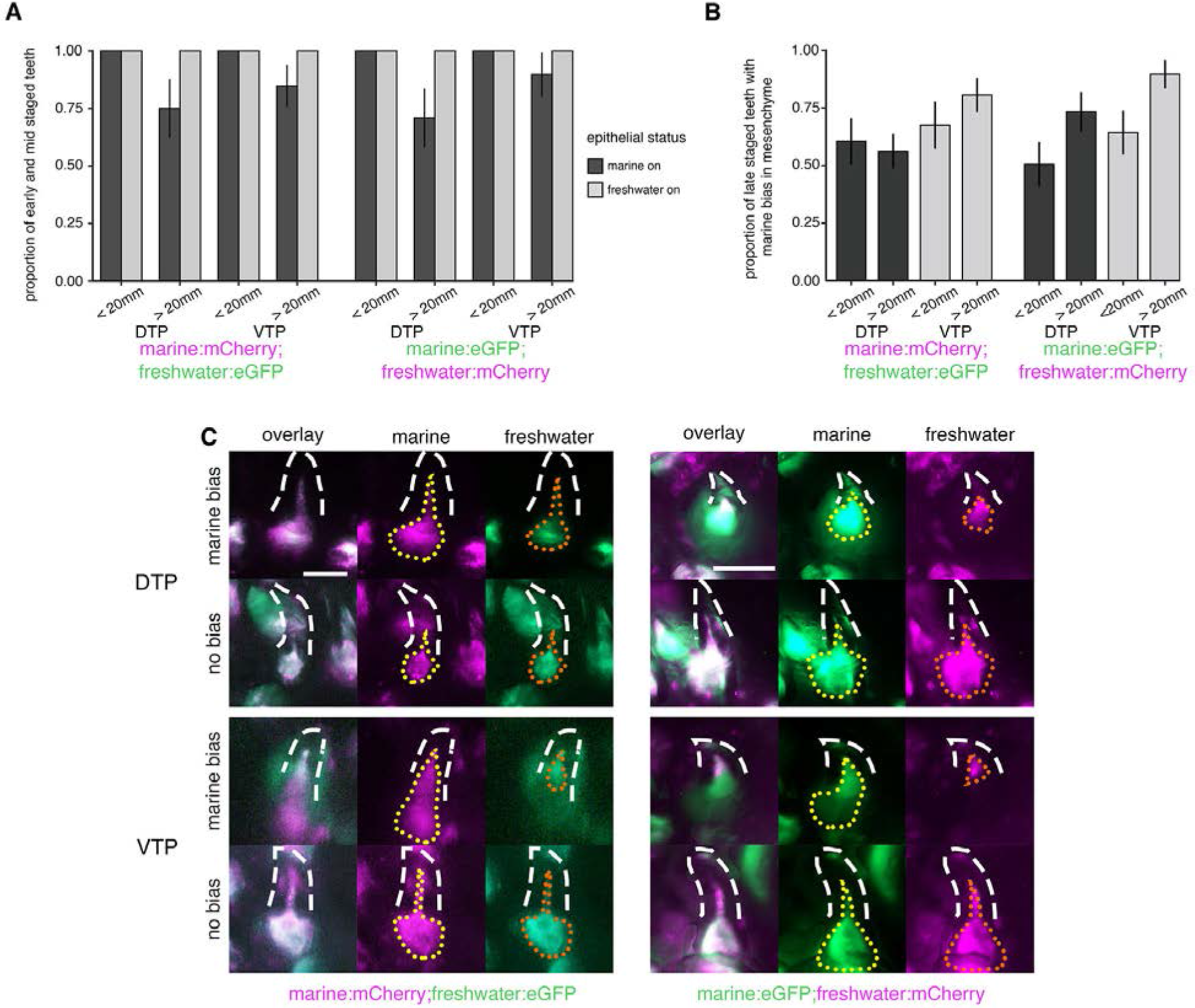
Differences in enhancer activity vary based on dorsal vs. ventral tooth field, fish total length, and epithelial vs. mesenchymal domain. (**A**) In < 20mm total length (pre-tooth number divergence) fish, the marine and freshwater alleles were expressed in the epithelium of all developing tooth germs regardless of genotype, while in > 20 mm total length (post-tooth number divergence) fish epithelial expression differences were consistent across tooth plates and genotypes. The freshwater allele consistently drove expression in all tooth germs scored, while the marine allele did not. (**B**) The proportion of erupted teeth that demonstrated an observed mesenchymal bias of an expanded marine enhancer domain differed across dorsal and ventral tooth plates (DTP and VTP, respectively), with more bias ventrally than dorsally. (**C**) Examples of erupted teeth (white dashed lines) from both DTP and VTP that were scored as either having a marine bias in the mesenchyme [if the freshwater enhancer mesenchymal domain (orange dotted line) was more restricted compared to the marine enhancer domain (yellow dotted line)], or no bias if the freshwater enhancer mesenchymal domain was equivalent to the marine enhancer domain. Scale bars = 50μm (**C**).

A bias towards the marine allele in the mesenchyme was observed in nearly every early or middle stage tooth germ, while the lack of bias, or entirely overlapping mesenchymal expression, was almost exclusively observed in late stage (erupted) tooth germs (Table S4). The ventral tooth plates had an increased prevalence of marine enhancer bias in the mesenchyme of individual teeth compared to the dorsal tooth plates (marine:mCherry;freshwater:eGFP ventral: 146/167 [87.4%], dorsal: 102/149, [68.5%] and marine:eGFP;freshwater:mCherry ventral: 123/136 [90.4%], dorsal: 122/149 [81.9%]). In early and middle stage teeth, we observed a consistent marine bias in the mesenchyme of both the ventral and dorsal tooth plates. In fully formed erupted teeth, a difference between the tooth plates became apparent. A larger proportion of erupted teeth were observed to have a marine bias in the mesenchyme in the ventral tooth plate compared to the dorsal tooth plate (Figure 5B-C).

There was a reduction in the proportion of erupted teeth with a marine bias when comparing post to pre divergence fish for all integrations and tooth plates (pre-divergence marine:mCherry;freshwater:eGFP ventral 54/80 [67.5%], dorsal 55/91 [60.4%] and marine:eGFP;freshwater:mCherry ventral 63/98 [64.3%], dorsal 51/103 [49.5%]) (Figure 5B) except for the dorsal tooth plates in the freshwater:eGFP;marine:mCherry genotype. Overall a bias towards marine expression in the mesenchyme was observed, with a consistently larger proportion of late stage teeth demonstrating a bias in the ventral teeth compared to the dorsal teeth, with the difference between tooth plates becoming more drastic in larger fish. Thus, the trend in marine mesenchymal bias across dorsal versus ventral tooth plates mirrors the chromosome 21 tooth number QTL, which had a 28 LOD greater effect on ventral pharyngeal tooth number than dorsal pharyngeal tooth number (Miller et al., 2014). In addition, the difference in bias between pre-divergence and post-divergence fish is consistent with allele specific expression data in which early in development the marine and freshwater alleles of *Bmp6* are expressed at more similar levels, while in older fish there is a *cis*-regulatory reduction in expression of the freshwater allele (Cleves et al., 2014).

### Pectoral and caudal fin expression differences

The *Bmp6* intron 4 enhancer was previously known to drive expression in the developing fin margins of the pectoral and caudal fins early in development, starting approximately 4 dpf (Cleves et al., 2018). In pre-hatching fish, 6 dpf, the domains of the two enhancers appear to be identical (Figure S4A). We found that enhancer activity persists at later stages in both the pectoral and caudal fins, specifically in the intersegmental joints. The fin rays of all fins in sticklebacks consist of a series of repeated segments, made up of hemi-segments encasing a mesenchymal core like other teleosts (Haas, 1962; Santamaría et al., 1992). In the caudal fin of both genotypes (freshwater:eGFP;marine:mCherry and freshwater:mCherry;marine:eGFP), the freshwater enhancer was observed to have activity in multiple intersegmental joints, while the activity of the marine enhancer was detected in none or few joints (Figure S4B). A similar pattern is observed in the pectoral fins (Figure S4C). With both enhancers, more basal joints were observed to have expression, while fluorophore intensity diminished as the joints became more distal. Overall, across both fin types, the freshwater allele appeared to be active in a larger number of intersegmental joints. While more proximal intersegmental joints were more likely to have activity from both enhancers, the most proximal joint was observed to be lacking detectable reporter expression in some fin rays (Figure S5A&B), suggesting a dynamic cycle of initial inactivity in newly formed, distal, intersegmental joints, followed by a period of activity in most joints as they adopt a more proximal identity, and a final transition to inactivity in the proximal most joints just prior to the ultimate fusion of basal most segment to the next segment.

### *Bmp6* expression differences between marine and freshwater fish

Given the consistent differences in reporter gene activity observed for the marine and freshwater enhancers, we next asked if endogenous *Bmp6* expression differed in tooth germs between marine and freshwater animals in a similar fashion. To answer this, we performed *in situ* hybridization (ISH) on thin sections of pharyngeal tissues from marine (Rabbit Slough) and freshwater (Paxton Benthic) adults (∼40 mm standard length). Marine and freshwater samples were collected, prepared, and assayed in parallel to ensure maximal comparability of the resulting data (see Methods). While early bud and cap stage tooth germs did not show any consistent differences in gene expression, we did observe more widespread mesenchymal expression in marine tooth germs at early and late bell stages, and consistently widespread IDE expression in freshwater epithelium relative at late bell stages (Figure 8). These ISH results corroborate the reporter construct activity, suggesting that the regulation of *Bmp6* mRNA in tooth germs varies in the same direction as the variation in activity seen between the marine and freshwater *Bmp6* intron 4 enhancers.

## DISCUSSION

### Freshwater and marine alleles of *Bmp6* tooth enhancer drive expression differences in developing teeth

Throughout the development of a tooth, multiple pathways and signals, including BMPs, are involved in organ initiation and growth. Knocking out the receptor *Bmpr1a* in the dental epithelium of mice leads to arrested development of the tooth at the bud stage, demonstrating a key activating role for BMP signaling during tooth development (Andl, 2004). Overexpressing *Noggin*, a BMP antagonist, in the epithelium also results in arrest at the placode stage (Wang et al., 2012). In addition, in *Msx1* mutant mice, exogenous *Bmp4* can rescue tooth development (Bei et al., 2000). Together, these results suggest a dynamic role of *Bmp* signaling in tooth development, both promoting and inhibiting tooth development at different stages. *Bmp6* is dynamically expressed during stickleback tooth development. Expression is detected early in the overlying inner dental epithelium (IDE) as well as in the condensing underlying odontogenic mesenchyme, with a subsequent cessation of expression in the epithelium, and continuous expression in the mesenchyme of the ossifying tooth (Cleves et al., 2014; Ellis et al., 2016). Freshwater sticklebacks homozygous for mutations in *Bmp6* have reductions in tooth number, showing *Bmp6* is required for aspects of tooth development in fish (Cleves et al., 2018).

A previously identified freshwater high-toothed associated haplotype within intron 4 of *Bmp6* underlies an evolved increase in tooth number. The core haplotype is defined by six polymorphic sites in the 468 bp region upstream of a minimally sufficient *Bmp6* tooth enhancer, potentially modifying enhancer activity. Three lines of evidence (the bicistronic line, and two line of reciprocal two-color lines) support the hypothesis that the associated polymorphisms upstream of the *Bmp6* tooth enhancer result in evolved spatial shifts in enhancer activity between the marine and freshwater alleles (Figures 1-4). Both alleles drove expression in the epithelium of early developing teeth, and in dental mesenchyme throughout development, similar to the expression pattern of the adjacent minimally sufficient 511 bp tooth enhancer previously reported (Cleves et al., 2018) as well as the reported expression of the endogenous *Bmp6* gene during tooth development (Cleves et al., 2014). In all three different transgenic comparisons, we observed the freshwater, high-toothed associated enhancer allele maintained a more robust expression domain in the overlying epithelium for a longer portion of a tooth’s development compared to the marine, low-toothed associated allele in multiple independent lines. Conversely, the marine allele appeared to drive reporter expression in a larger domain in the underlying mesenchyme in a large proportion of teeth. We additionally found that marine and freshwater endogenous *Bmp6* gene expression domains differed in a manner that was consistent with the reporter gene results. Specifically, we observed larger mesenchymal domains in marine relative to freshwater fish, and expanded IDE domains in freshwater relative to marine fish, especially in late bell stage tooth germs. Together these data support the hypothesis that the intron 4 enhancer variants associated with tooth number differences drive *Bmp6* expression differences in tooth germs of >20 mm fish, which in turn leads to evolved tooth gain in freshwater fish (Figure 7). Outstanding questions include what these deep mesenchymal cells are and whether the expanded marine mesenchymal domain might include quiescent mesenchymal cells involved in tooth replacement. Other important questions include whether the differential expression of the endogenous *Bmp6* gene that occurs between marine and freshwater fish is at least partially driven by the two enhancer alleles, and if so, what the allelic effects are on tooth development and replacement, and which mutations are responsible for the expression differences.

**Figure 6.**
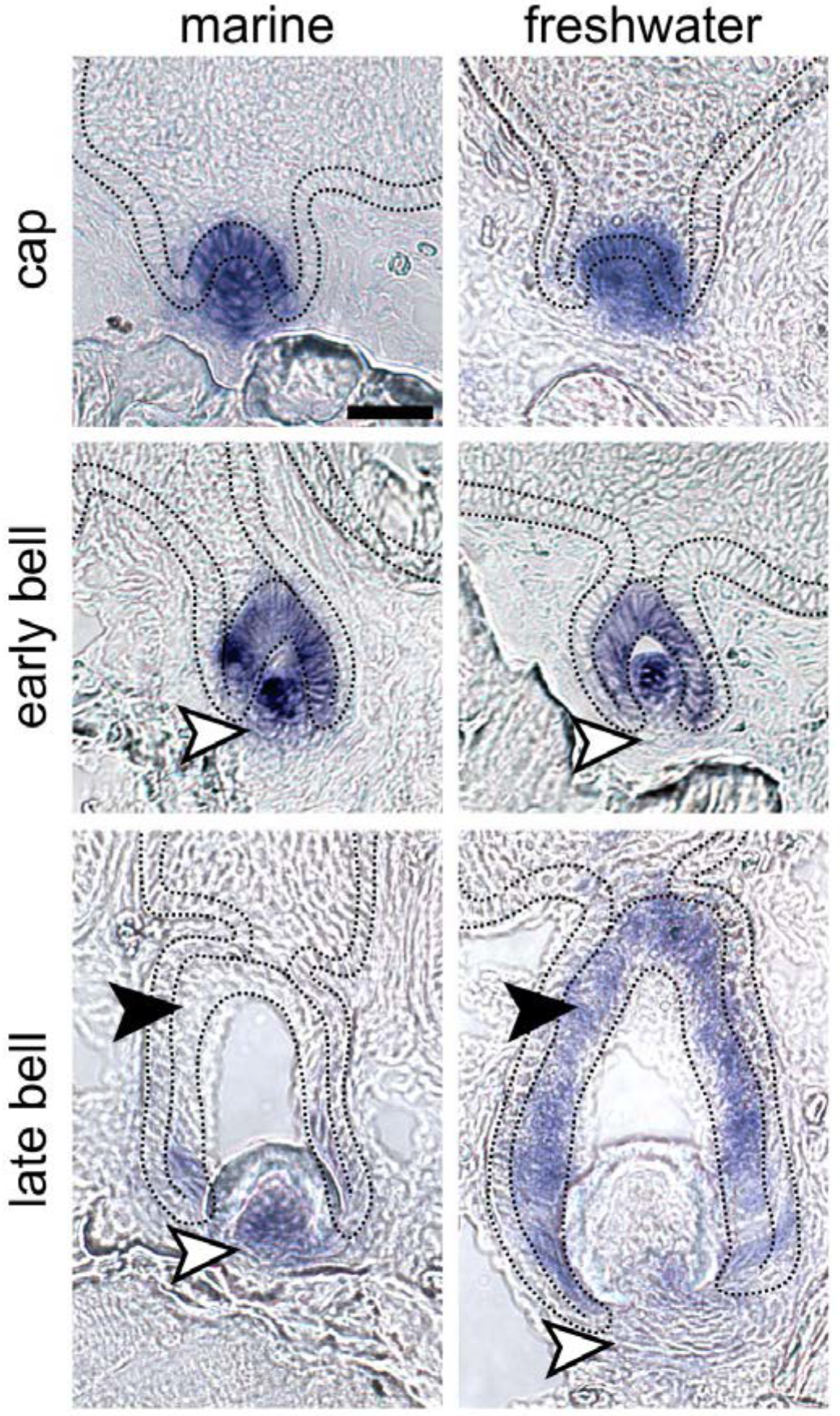
*In situ* hybridization illustrates that *Bmp6* expression shifts mirror enhancer activity differences in marine and freshwater backgrounds. *In situ* hybridization (ISH) of *Bmp6* expression on thin sections of marine (left column) and freshwater (right column) homozygous backgrounds suggest that marine fish exhibit expanded mesenchymal expression at early and late bell stages (white arrowheads in middle and bottom rows, respectively), while freshwater fish exhibit relatively broader expression in the inner dental epithelium (IDE) of late bell stage teeth (black arrowheads in bottom row). No expression domain differences were observed in cap stage tooth germs (top row). Marine and freshwater strains are derived from population in Rabbit Slough, AK, USA (RABS), and Paxton Lake, BC, Canada (PAXB), respectively. Black dotted lines demarcate the basalmost layer of epithelium, adjacent to the basement membrane, which includes the inner and outer dental epithelium. See Figure S6 for DAPI counterstains and ISH images without markup. Scale bar = 20μm and applies to all panels.

**Figure 7.**
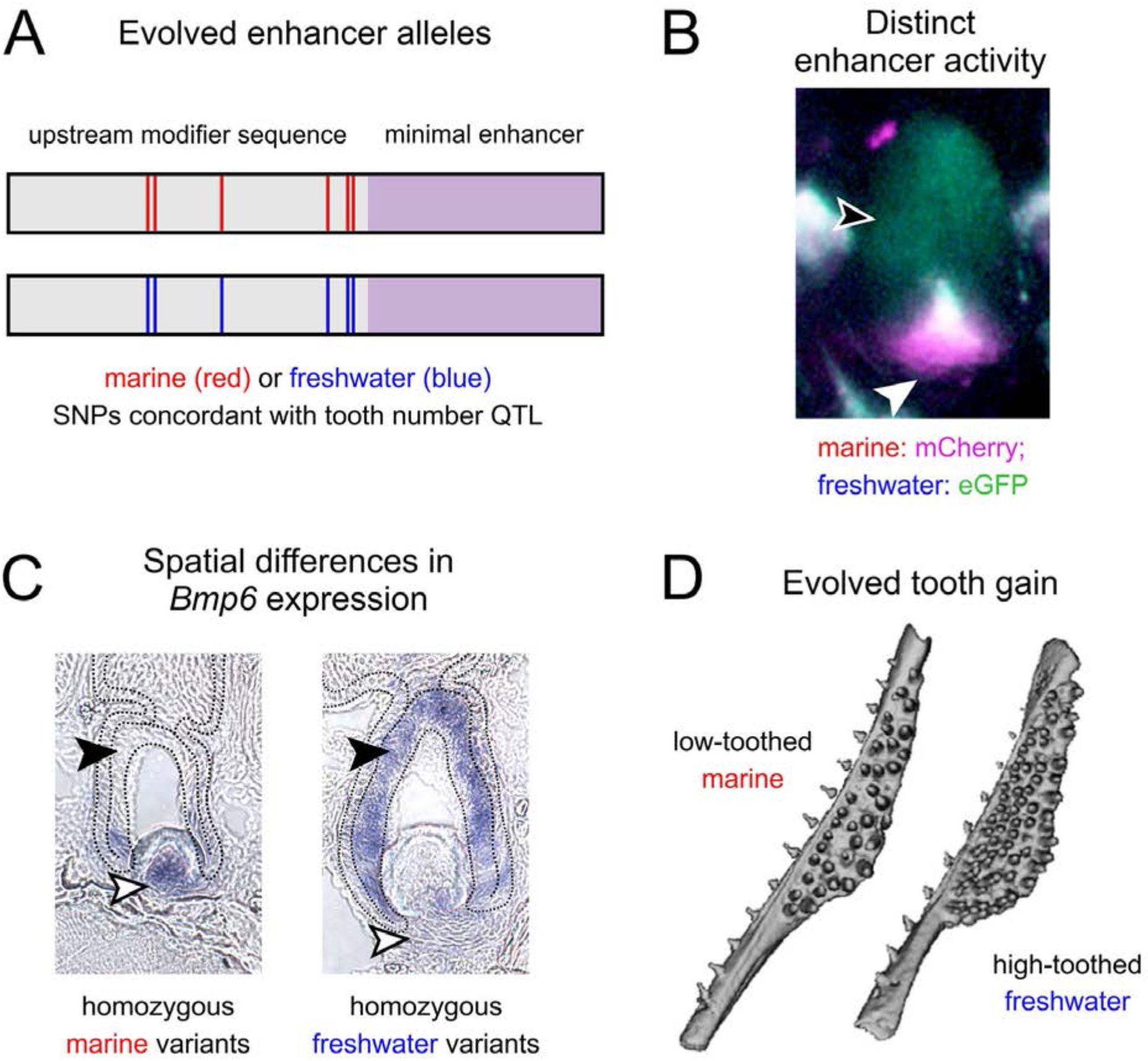
A model for the role of *Bmp6 cis* regulatory changes in underlying evolved tooth gain in sticklebacks. **(A)** Quantitative trait loci (QTL) and fine mapping previously revealed variants in intron 4 of *Bmp6* that were associated with evolved tooth gain in freshwater fish (Cleves et al., 2014, 2018; Miller et al., 2014). These variants are adjacent to a previously characterized minimal enhancer (lavender) that was shown to drive expression in tooth epithelium and mesenchyme (Cleves et al., 2018). Six core single nucleotide polymorphisms (SNPs, depicted as red and blue lines within the modifier sequence), showed complete concordance with a large effect tooth number QTL (Cleves et al., 2018). **(B)** Marine and freshwater enhancers have different spatial activity, with the derived freshwater allele driving less mesenchymal expression, but more epithelial expression relative to the marine allele. **(C)** Consistent with the different enhancer activity, *Bmp6* expression by *in situ* hybridization is reduced in the mesenchyme but expanded in the epithelium in freshwater teeth relative to marine teeth. **(D)** We hypothesize that the enhancer alleles (A) have spatially shifted enhancer activity (B), resulting in shifts in *Bmp6* expression overall (C), and evolved tooth gain in freshwater fish (D).

Previous allele specific expression (ASE) experiments demonstrated a 1.4 fold reduction in the freshwater *Bmp6* allele compared to the marine in F_1_ hybrid adult tooth tissue that included the entire ventral pharyngeal jaw, and thus both tooth epithelial and mesenchymal cells (Cleves et al., 2014). The mesenchymal biases in reporter expression are consistent with the ASE result, with more robust mesenchymal expression driven by the marine allele compared to the freshwater allele potentially responsible for the higher expression of the marine allele in the ASE experiments. In contrast, the expanded freshwater epithelial enhancer domain is not consistent with the overall ASE result in which freshwater alleles had *cis*-regulatory downregulation relative to marine alleles. Since the reduced mesenchymal domain in the freshwater enhancer relative to the marine enhancer was the most striking qualitative difference, it is possible that the epithelial bias, with a stronger signal driven by the freshwater enhancer, is quantitatively canceled out by the bias in the mesenchyme, explaining the overall reduction of freshwater *Bmp6* expression compared to marine *Bmp6* expression in F_1_ hybrids.

The enhancer expression differences appeared more pronounced in larger, post tooth number divergence fish compared to smaller, pre tooth number divergence fish. While the mesenchyme appeared to have a somewhat reduced difference of expression between the two alleles, the epithelium demonstrated less pronounced differences in activity between the alleles in pre-divergence fish. The observation is consistent with ASE results and the divergence in tooth number in marine and freshwater fish. While the mesenchymal difference was still observable early, it is possible there are other regulatory regions which act as repressors for the marine *Bmp6* allele or enhancers for the freshwater *Bmp6* allele early in development and so mask the mesenchymal bias of the marine intron 4 enhancer. For example, we previously reported a 5’ *Bmp6* tooth enhancer that likely also contributes to the overall pattern of *Bmp6* in developing teeth (Erickson et al., 2015).

Future experiments to measure ASE in isolated tissues, with epithelium and mesenchyme separated could test whether opposing quantitative differences are present in dental epithelium vs. mesenchyme, as the new data presented here suggest. A quantitative method could be used to further test a hypothesis in which the two enhancers drive differing levels of expression, such as pyrosequencing (Wittkopp, 2012) with the two enhancers both driving identical fluorophores, with a single synonymous mutation distinguishing the two. Alternatively, single-cell RNA-seq (scRNA-seq) in the dental epithelium and mesenchyme, targeting the respective reporters of each enhancer, could determine if there are quantifiable expression differences between the two enhancers.

### QTL-associated sequence difference in alleles may underlie expression domain differences

There are 14 point mutations and three indels distinguishing a low-toothed marine (Little Campbell) allele from the high-toothed Paxton Lake allele of the intron 4 enhancer in our reporter constructs. Previous experiments identified ten of these SNPs that co-occur consistently with the presence or absence of a tooth number QTL and of these ten, the core six are present in the enhancer reporter constructs tested here (Cleves et al., 2018). From our results we are unable to distinguish whether these six polymorphisms contribute to the expression differences we observed. While it is possible that the three indels or the eight non-QTL-associated SNPs may contribute, it is an attractive and parsimonious hypothesis that the same SNPs that co-occur with the tooth QTL are also responsible for the reporter expression differences, and the previously described allele specific expression results. Of the six QTL-associated SNPs tested here, of special interest is the second QTL-associated SNP (CßàT), which in the freshwater allele, creates a predicted NFATc1 binding site (Cleves et al., 2018). NFATc1 was shown to have importance in the balancing of quiescent and actively dividing stem cells in hair follicles (Horsley et al., 2008) which share homology with teeth (Ahn, 2015; Biggs & Mikkola, 2014; Pispa & Thesleff, 2003), and so a difference in NFATc1 binding may potentially play a role in the *Bmp6* allele specific expression and enhancer activity differences observed previously and here. Supporting this hypothesis, *Nfatc1b* expression was recently shown to be present in stickleback tooth germs and functional tooth mesenchyme (Square et al., 2021).

To better determine which polymorphisms may underlie the expression differences we observed, hybrid enhancers can be made. For example, if the creation of an NFATc1 binding site is at least partially responsible for the observed differences, a marine allele with the SNP converted to the freshwater identity, from a ‘C’ to a ‘T’, may recapitulate the freshwater enhancer expression patterns. By creating and testing hybrid enhancers, future experiments could test which enhancer polymorphisms alone and in combination, contribute to the expression differences reported here.

### Fin expression differences

In addition to the reporter expression differences driven by the two enhancers during tooth development, we observed distinct expression patterns in the pectoral and caudal fins. It was previously known that the minimal 511 base pair enhancer drove expression in the margins of early pectoral and median fins, but expression in adult fins had not been described. BMP signaling plays a role in fin regeneration, with BMP inhibition reducing osteoblast differentiation in new cells arising at the leading edge of the regenerating fin (Stewart et al., 2014). During zebrafish fin regeneration *bmp2b, bmp4,* and *bmp6* are expressed, and are thought to be important (Laforest et al., 1998; Murciano et al., 2002; Quint et al., 2002; Smith et al., 2006). While both alleles of the *Bmp6* enhancer drive expression in the pectoral and caudal fins of sticklebacks, the differing enhancer activities may result in developmental differences, through osteoblast function in the developing lepidotrichia and intersegmental joints, possibly leading to different fin morphologies and/or regenerative abilities. Differences in expression of *bmp2* have been observed in the regeneration of different rays of the caudal fin in cichlids (Ahi et al., 2017), as well as the expression of the gene *msxb*, which is downstream of *bmp* signaling in the regenerating zebrafish fin (Smith et al., 2006).

Multiple studies have identified habitat specific differences in fin morphology (Hendry et al., 2011; Kristjánsson et al., 2005; Taylor & McPhail, 1986). As the two enhancers are derived from populations with two distinct ecotypes, a benthic freshwater population, and a highly mobile anadromous population, it is possible this enhancer may influence pectoral and caudal fin size and shape in an adaptive manner. Characterization of fin morphology using fish from either a population in which the high-toothed and low-toothed associated haplotypes are segregating, or those from a control cross in which both alleles were present in the founding, could test whether there is a fin morphology difference associated with the different alleles.

### Bicistronic constructs and the use of genetic insulators

Simultaneous comparison of two enhancer alleles in a single organism via a bicistronic construct is an attractive means to compare molecularly divergent enhancers (e.g. pairs of enhancers that contain sequence variation across populations to determine if there are population specific differences of enhancer activity). Previous work in zebrafish utilized genetic insulators as part of an enhancer trap as well as with two different tissue specific promoters and demonstrated the effectiveness of the technique (Bessa et al., 2009; Shimizu & Shimizu, 2013).

Here we used a bicistronic construct with a *Bmp6* enhancer and *Col2a1a* enhancer driving different fluorophores in mosaically transgenic F_0_ fish to test whether the activities of two enhancers could be insulated from each other. Within the same F_0_ individual, some domains demonstrated a high degree of insulator effectiveness while others did not. There are at least two possible explanations: 1) the insulated vs. non-insulated regions represent distinct and mosaic integration events, with the insulator effectiveness determined by the integration site in a particular subpopulation of cells, or 2) the same integration event can differ in insulator behavior stochastically or based on some context that differs from an insulated expression domain to an un-insulated domain. Regardless, examining enhancer activity in stable lines will still provide a more complete picture of the role of the regulatory element and has advantages over mosaic F_0_ analyses.

Genetic insulators have been reported to limit enhancer activity across the insulator boundary (Bessa et al., 2009; Shimizu & Shimizu, 2013) as well as protect against position effects (Chung et al., 1993), while other experiments show a lack of protection (Grajevskaja et al., 2013). The insulator used here, from the 5’end of the mouse tyrosinase locus, was reported to bind CTCF, like the β-globin 5’HS4 insulator from chicken, and is reported to prevent influences from nearby chromatin state and gene activity, the hallmarks of genetic insulators (Giraldo, 2003; Molto et al., 2009; Montoliu et al., 1996). As there are conflicting reports of the use of insulators to fully shield from nearby chromatin states and position effects, the combined use of a landing pad locus could help to further reduce these effects (Roberts et al., 2014). We recommend a multiple pronged approach utilizing multiple transgenic lines (e.g. either bicistronic constructs or multiple independent reciprocal two-color lines where each enhancer drives a different fluorophore in the same animal). Similar methods in doubly transgenic animals should allow future dissection of spatial differences in enhancer alleles, with the two methods acting as means of independent verification.

Changes in *cis*-regulation of developmental genes can be an important driver of morphological evolution, as well as human disease. The impact of mutations in *cis*-regulatory regions can be difficult to predict, and if the effect is subtle or slight, also to detect. The use of two enhancers in the same individual, either as parts of two independent transgenes or within a single bicistronic construct, can both control for the trans-environment and make even slight differences in expression activity apparent due to simultaneous imaging of reporter genes driven by both enhancers. Such an approach allows for directly comparing molecularly divergent regulatory elements, potentially identifying causal polymorphisms with important developmental and evolutionary implications.

## DATA AVAILABILITY STATEMENT

Strains and plasmids are available upon request. The authors affirm that all data necessary for confirming the conclusions of the article are present within the article, figures, and tables.

## ACKNOWLEDGEMENTS

We thank Phillip Cleves, James Hart, Priscilla Erickson, Ana Shaughnessy for helpful discussions, and Sophie Archambeault and Alyssa Borman for comments on the manuscript.

## Supplemental materials

### Supplemental figures

**Figure S1.**
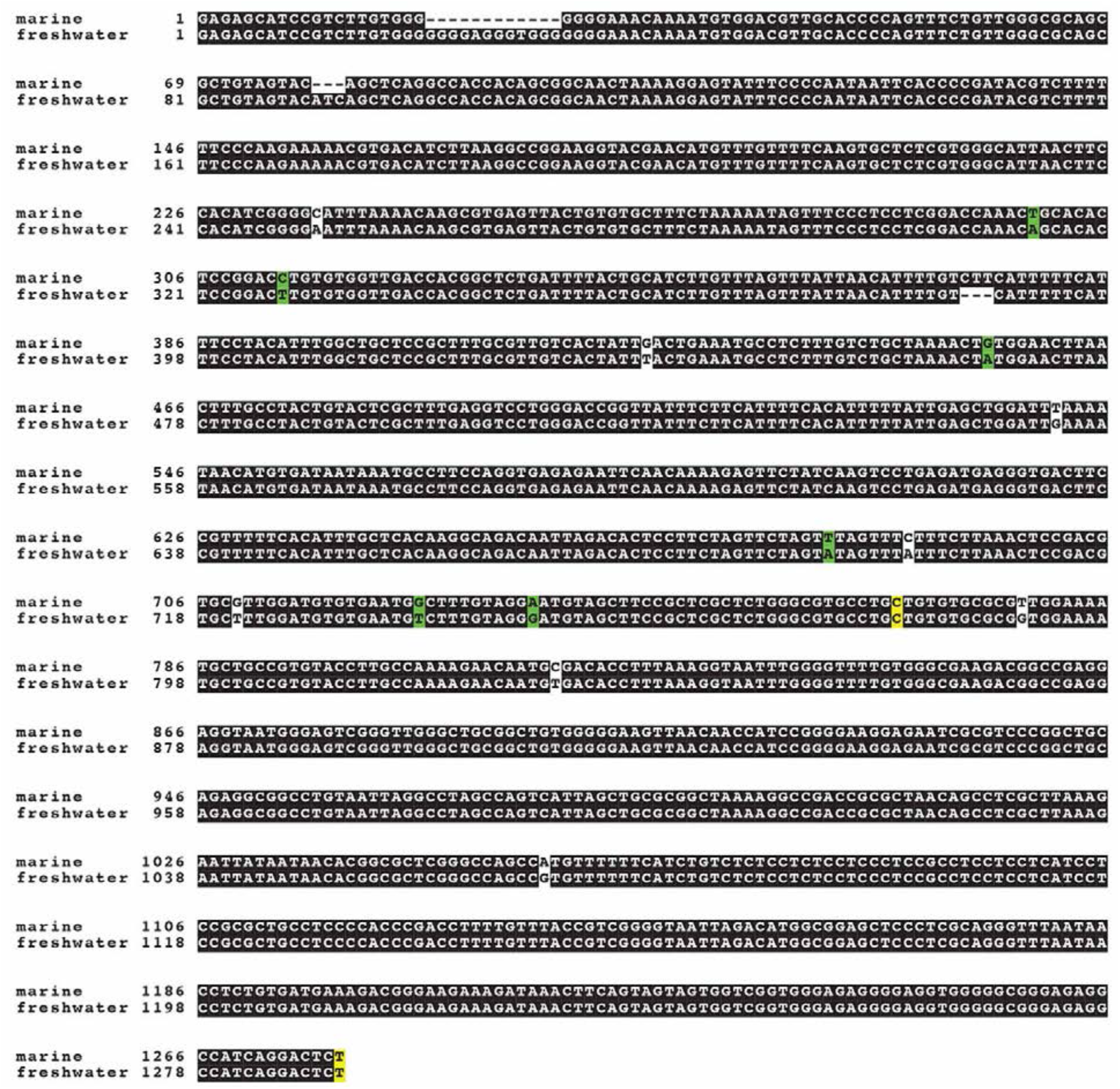
Sequence alignment of marine and freshwater alleles of *Bmp6* tooth enhancer. Six core single nucleotide polymorphisms (green) concordant with the presence or absence of a large effect tooth number QTL lie upstream of a ∼511 bp minimal *Bmp6* tooth enhancer (start and end in yellow). Other polymorphisms (white) are not concordant with the presence or absence of the tooth QTL (Cleves et al., 2018).

**Figure S2.**
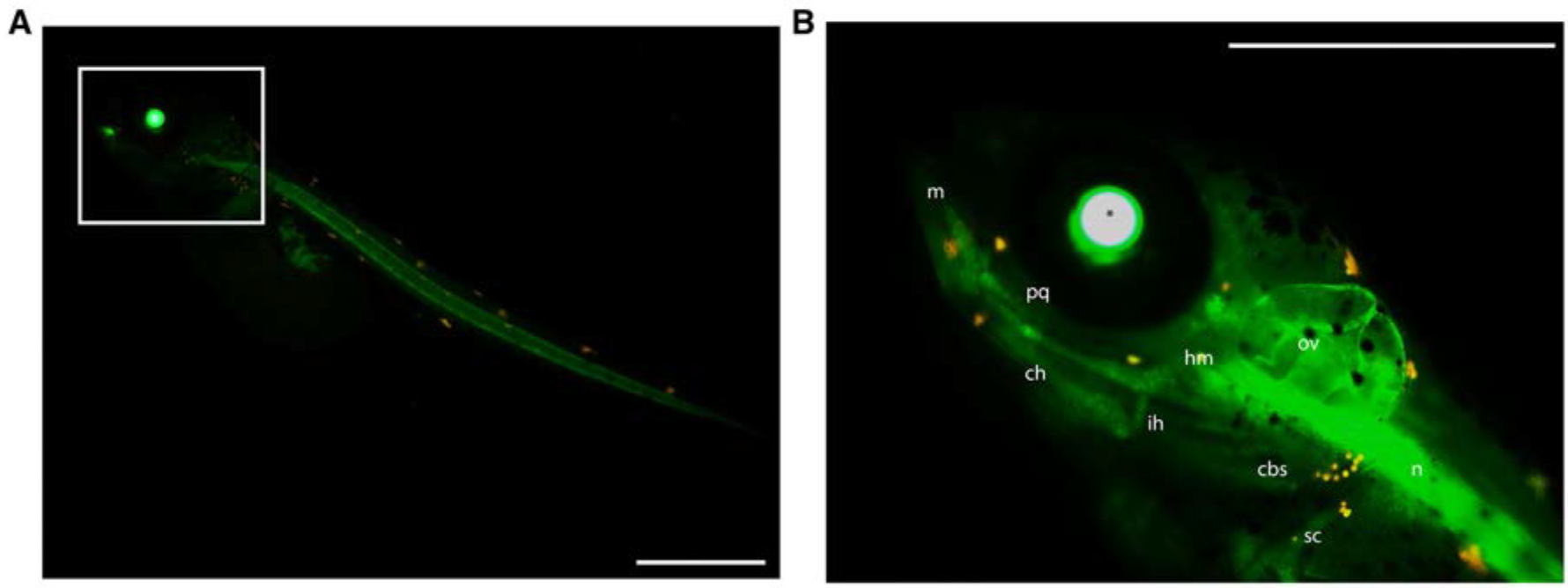
A Col2a1 enhancer drives reporter expression in craniofacial cartilage and notochord in developing stickleback embryos. (**A**) In a ten day post-fertilization embryo, reporter expression was observed in notochord (n) and (**B**) craniofacial cartilage including Meckel’s (m), ceratohyal (ch), interhyal (ih), ceratobranchials (cbs), palatoquadrate (pq), and the hyosympletic (hm). Expression was also seen in the scapulocoracoid (sc), and otic vesicle (ov). Scale bars = 500μm.

**Figure S3.**
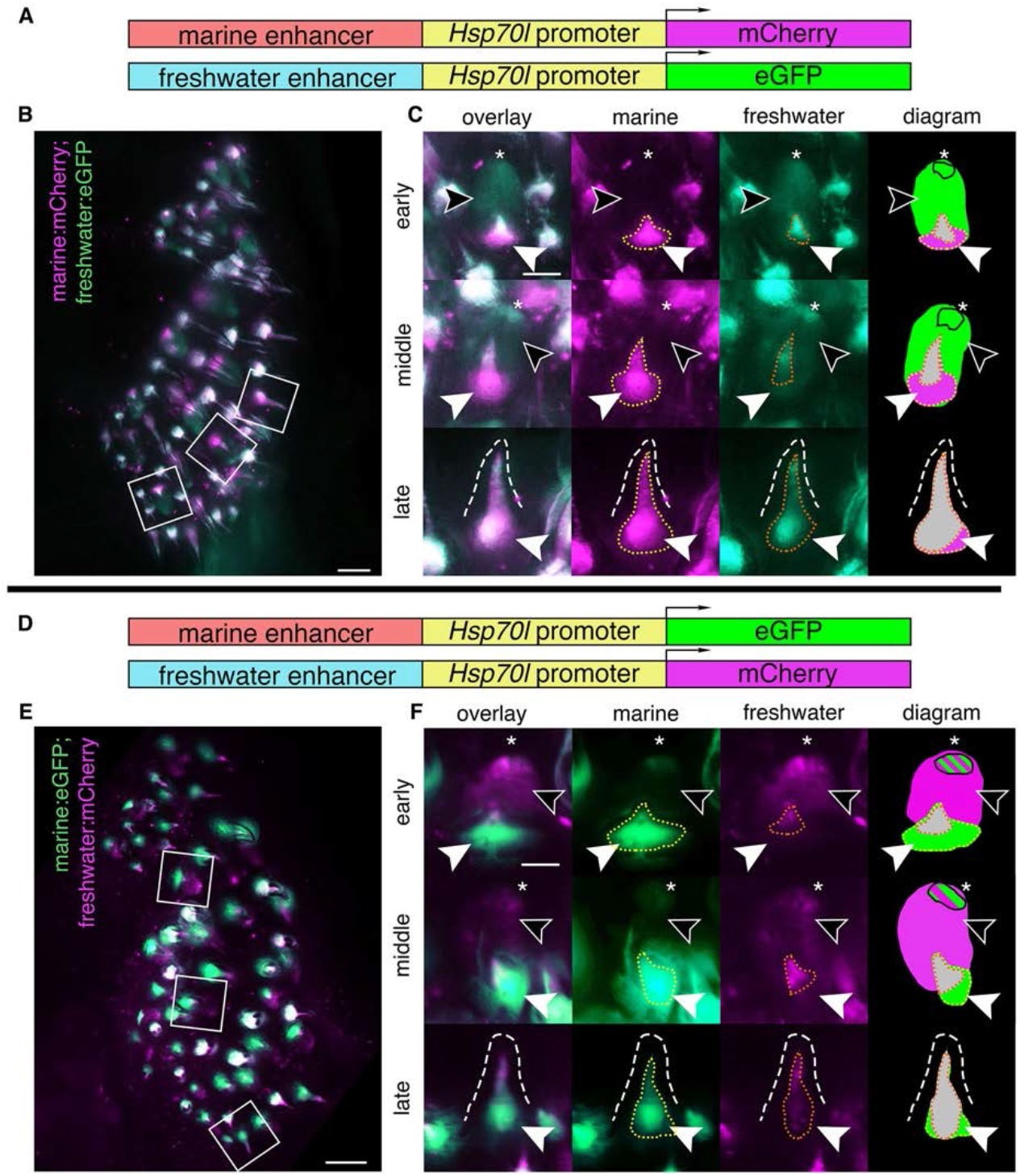
Marine and freshwater *Bmp6* enhancers drive different spatial patterns in dorsal pharyngeal teeth. Dorsal pharyngeal tooth plates from fish doubly transgenic for two alleles of the *Bmp6* intron 4 enhancer driving two different reporter genes (**A,D**): the marine enhancer driving mCherry with the freshwater enhancer driving eGFP (**B,C**) and the marine enhancer driving eGFP with the freshwater enhancing driving mCherry (**E,F**). Unilateral dorsal pharyngeal tooth plates (**B,E**) are shown, next to representative teeth from three stages (**C,F**): early, middle, and late highlighted by white boxes in **B,E**. (**C,F**) Early: freshwater enhancer drove expression in the epithelium (black arrowheads), with concentrated expression in the tip (asterisk), while the marine enhancer did not reliably drive expression in the epithelium, but was observed in the distal tip (**F**) in some instances. Both enhancers also drove expression in the mesenchyme (solid white arrowhead) with a larger expression domain of the marine allele (yellow dotted line) compared to the freshwater allele (orange dotted line). Middle: freshwater allele still drove expression in the epithelium while the marine allele was restricted to the distal tip. The marine allele drove more robust mesenchymal expression compared to the freshwater allele. Late: marine allele drives robust expression in the mesenchyme compared to freshwater allele in mineralized tooth (dashed line). Diagram: summary of tooth epithelial and mesenchymal domains. The relative sizes of green and magenta hatched lines correspond to the approximate relative strength of expression in the epithelium. Overlapping mesenchyme domain is grey, and expanded marine mesenchyme is marked with white arrowhead. Scale bars = 100μm (**B**,**E)**, 50μm (**C**, **F)**.

**Figure S4.**
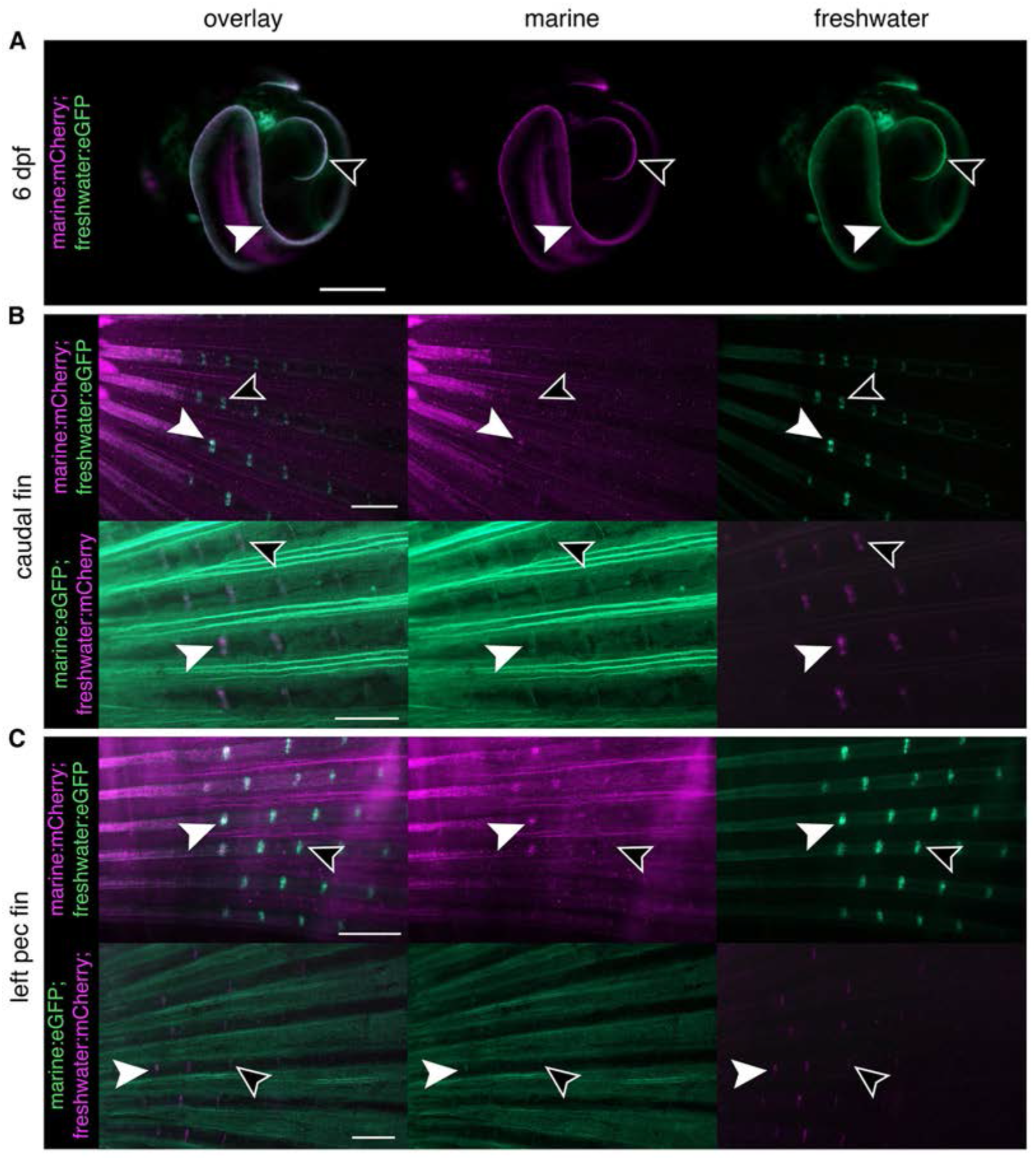
Freshwater allele drives expression in more intersegmental joints of both pectoral and caudal fins compared to the marine allele. (**A**) In young, pre-hatching fish (6 dpf) the marine and freshwater enhancers drive expression in identical patterns in the developing fin margins of the pectoral fins (solid white arrowhead) and median fin (black arrowhead). (**B**) In adult caudal fins the more basal intersegmental joints were observed to have activity from both the marine and freshwater alleles (solid white arrowhead) while more distal joints were observed to only have freshwater enhancer activity (black arrowhead). The pattern was observed across both enhancer/reporter pairings. (**C**) Left pectoral fins from adults were observed to have activity from both enhancers in more basal intersegmental joints (solid white arrowheads) while only the freshwater allele was observed to have activity in more distal joints (empty arrowheads). Scale bars = 0.5 mm.

**Figure S5.**
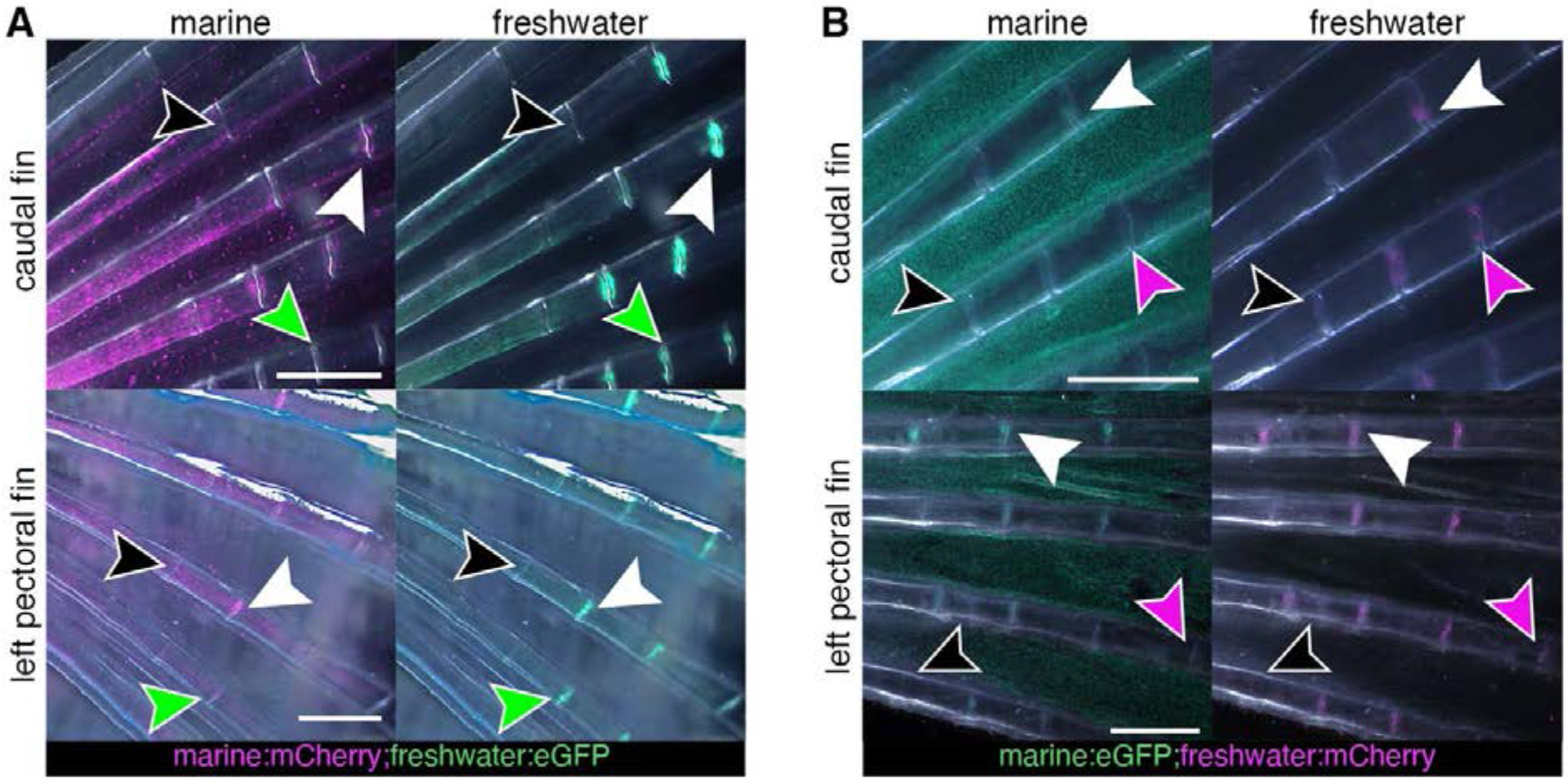
Fin expression patterns of both alleles change over developmental time. (**A)** Caudal and pectoral fins with the freshwater enhancer driving eGFP and marine enhancer driving mCherry. Only the freshwater enhancer is active in more distal joints (green arrowhead) while in more basal joints both enhancers are active (solid white arrowhead). No enhancer activity was observed in the most basal joints (black arrowhead). (**B**) Caudal and pectoral fins with the freshwater enhancer driving mCherry and marine enhancer driving eGFP. Similar to (**A**), the freshwater allele is active in more distal joints than the marine allele (purple arrowhead), more basal joints exhibit activity from both enhancers (solid white arrowhead). In the most basal joints, activity from either enhancer was not observed (black arrowhead). Scale bars = 0.5mm.

**Figure S6.**
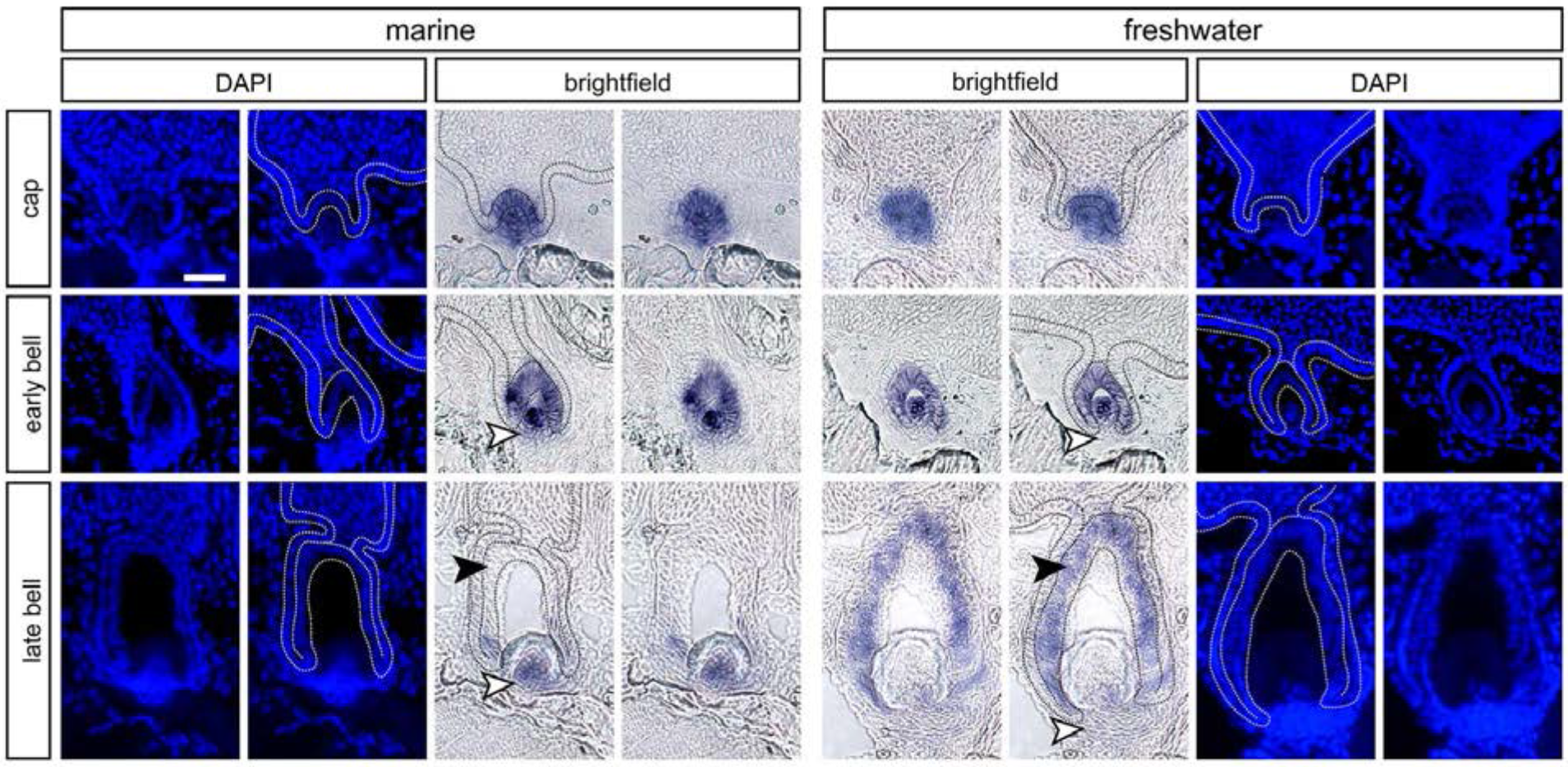
DAPI counterstain distinguishes between epithelial and mesenchymal tissues on thin sections. Inner four columns show brightfield *in situ* hybridization (ISH) images for *Bmp6* expression on marine (left) and freshwater (right) backgrounds, innermost columns with no annotations, adjacent to the same images with annotations (as presented in Figure 6). The outermost four columns show DAPI counterstains of the same sections, again shown both with and without annotations. The first row shows a cap stage tooth, the second row shows an early bell stage tooth, and the third row shows a late bell stage tooth. All dotted lines (black in brightfield images, white in DAPAI images) demarcate the basalmost layer of epithelium in the tooth field, which is contiguous with the inner and outer dental epithelia of tooth germs. Regions where differences in expression were detected are marked with arrowheads: white arrowheads mark expanded mesenchymal expression in marine relative to freshwater, while black arrowheads mark expanded epithelial expression in freshwater relative to marine (as shown in Figure 8). Scale bar = 20 μm and applies to all panels.

### Supplemental tables

**Table S1.** Insulator scores for bicistronic Col2a1a:mCh;Bmp6 tooth enhancer:eGFP transgene.

| Domain | “0”- apparent no insulation | “1” – partial insulation observed | “2”- apparent complete insulation | Total fluorescence positive domains |
| --- | --- | --- | --- | --- |
| Left pec fin | 24 | 13 | 28 | 65 |
| Right pec fin | 28 | 14 | 21 | 63 |
| Median fin | 34 | 23 | 29 | 86 |
| Notochord | 9 | 1 | 3 | 13 |
| Total | 95 | 51 | 81 | 227 |

For each reporter positive domain in F<sub>0</sub> fish with *Col2a1a*:mCh;*Bmp6* tooth enhancer:eGFP transgene, a score of 0-2 was given for observed non, partial, or complete insulation.

**Table S2.** Insulator scores for bicistronic Col2a1a:eGFP;Bmp6 tooth enhancer:mCh transgene.

| Domain | “0”- apparent no insulation | “1” – partial insulation observed | “2”- apparent complete insulation | Total fluorescence positive domains |
| --- | --- | --- | --- | --- |
| Left pec fin | 12 | 2 | 4 | 18 |
| Right pec fin | 6 | 4 | 3 | 13 |
| Median fin | 15 | 4 | 3 | 22 |
| Notochord | 5 | 0 | 5 | 10 |
| Total | 38 | 10 | 15 | 63 |

For each reporter positive domain in F<sub>0</sub> fish with *Col2a1a*:mCh;*Bmp6* tooth enhancer:eGFP transgene, a score of 0-2 was given for observed non, partial, or complete insulation.

**Table S3.**
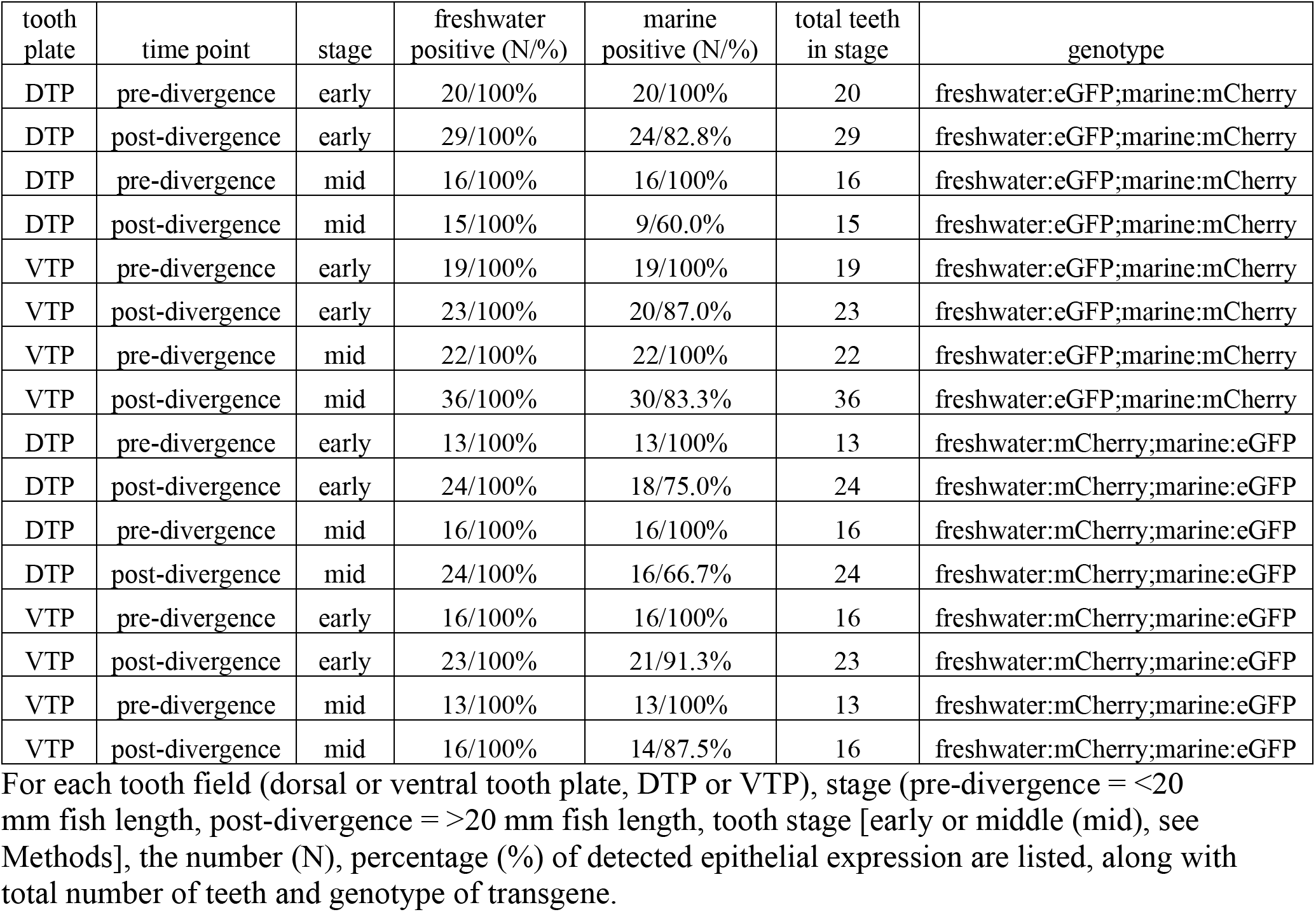
Epithelial expression of enhancer by tooth plate, tooth stage, and genotype.

**Table S4.** Mesenchymal bias of enhancer expression by tooth plate, tooth stage, and genotype.

| tooth plate | time point | stage | unbiased mesenchymal expression (N/%) | biased mesenchymal expression (N/%) | Total teeth in stage | genotype |
| --- | --- | --- | --- | --- | --- | --- |
| DTP | pre-divergence | early | 3/15% | 17/85% | 20 | freshwater:eGFP;marine:mCherry |
| DTP | post-divergence | early | 1/3.4% | 28/96.6% | 29 | freshwater:eGFP;marine:mCherry |
| DTP | pre-divergence | mid | 2/12.5% | 14/87.5% | 16 | freshwater:eGFP;marine:mCherry |
| DTP | post-divergence | mid | 0/0% | 15/100% | 15 | freshwater:eGFP;marine:mCherry |
| DTP | pre-divergence | late | 36/39.6% | 55/60.4% | 91 | freshwater:eGFP;marine:mCherry |
| DTP | post-divergence | late | 46/43.8% | 59/56.2% | 105 | freshwater:eGFP;marine:mCherry |
| VTP | pre-divergence | early | 4/21.1% | 15/88.9% | 19 | freshwater:eGFP;marine:mCherry |
| VTP | post-divergence | early | 0/0% | 23/100% | 23 | freshwater:eGFP;marine:mCherry |
| VTP | pre-divergence | mid | 2/9.1% | 20/90.9% | 22 | freshwater:eGFP;marine:mCherry |
| VTP | post-divergence | mid | 0/0% | 36/100% | 36 | freshwater:eGFP;marine:mCherry |
| VTP | pre-divergence | late | 26/32.5% | 54/67.5% | 80 | freshwater:eGFP;marine:mCherry |
| VTP | post-divergence | late | 21/19.4% | 87/80.6% | 108 | freshwater:eGFP;marine:mCherry |
| DTP | pre-divergence | early | 0/0% | 13/100% | 13 | freshwater:mCherry;marine:eGFP |
| DTP | post-divergence | early | 0/0% | 24/100% | 24 | freshwater:mCherry;marine:eGFP |
| DTP | pre-divergence | mid | 1/6.3% | 15/93.7% | 16 | freshwater:mCherry;marine:eGFP |
| DTP | post-divergence | mid | 0/0% | 24/100% | 24 | freshwater:mCherry;marine:eGFP |
| DTP | pre-divergence | late | 51/49.5% | 52/50.5% | 103 | freshwater:mCherry;marine:eGFP |
| DTP | post-divergence | late | 27/26.7% | 74/73.3% | 101 | freshwater:mCherry;marine:eGFP |
| VTP | pre-divergence | early | 0/0% | 16/100% | 16 | freshwater:mCherry;marine:eGFP |
| VTP | post-divergence | early | 0/0% | 23 (2 Freshwater [8.7%], 21 Marine [91.3%]) | 23 | freshwater:mCherry;marine:eGFP |
| VTP | pre-divergence | mid | 0/0% | 13/100% | 13 | freshwater:mCherry;marine:eGFP |
| VTP | post-divergence | mid | 0/0% | 16/100% | 16 | freshwater:mCherry;marine:eGFP |
| VTP | pre-divergence | late | 35/35.7% | 63/64.3% | 98 | freshwater:mCherry;marine:eGFP |
| VTP | post-divergence | late | 10/10.3% | 87 (1 Freshwater [1%], 86 [88.7%] Marine) | 97 | freshwater:mCherry;marine:eGFP |

For each tooth field (dorsal or ventral tooth plate, DTP or VTP), stage (pre-divergence = <20 mm fish length, post-divergence = >20 mm fish length, tooth stage [early or middle (mid), see Methods], the number (N), percentage (%) of detected mesenchymal bias in expression are listed, along with total number of teeth and genotype of transgene.

### Supplemental methods

Multiple fluorescent reporter transgenes were assembled using the methods and primers as described below. Component abbreviations below are as follows: *Hsp70l* = stickleback *Hsp70l* promoter (O’Brown et al., 2015); GAB = mouse tyrosinase insulator (Bessa et al., 2009); *Col2a1a* = *Col2a1a R2* enhancer (Dale & Topczewski, 2011).

#### *Col2a1a* containing insulator construct #1

##### *Col2a1a* enhancer/*Hsp70l*àmCh+GAB+eGFPß*Hsp70l*/*Bmp6* enhancer

The components of GAB, eGFP, and *Hsp70l*/*Bmp6* enhancer were amplified using primers MDS126/136, MDS137/89, and MDS90/131 respectively. The amplicons were combined with a modified plasmid (pT2He, modified to contain only polyclonal sites) linearized with *Nde*I and *BamH*I as well as Gibson Assembly master mix (NEB #E2611L) and incubated following the manufacturer’s protocol. The resulting plasmid was digested with *Nde*I and *Bsu36*I and the fragments for the second half, *Col2a1a* enhancer/*Hsp70l* and mCherry, were amplified with MDS138/139 and MDS140/141 respectively. The plasmid and amplicons were combined with Gibson Assembly master mix and incubated following the manufacturer’s protocol.

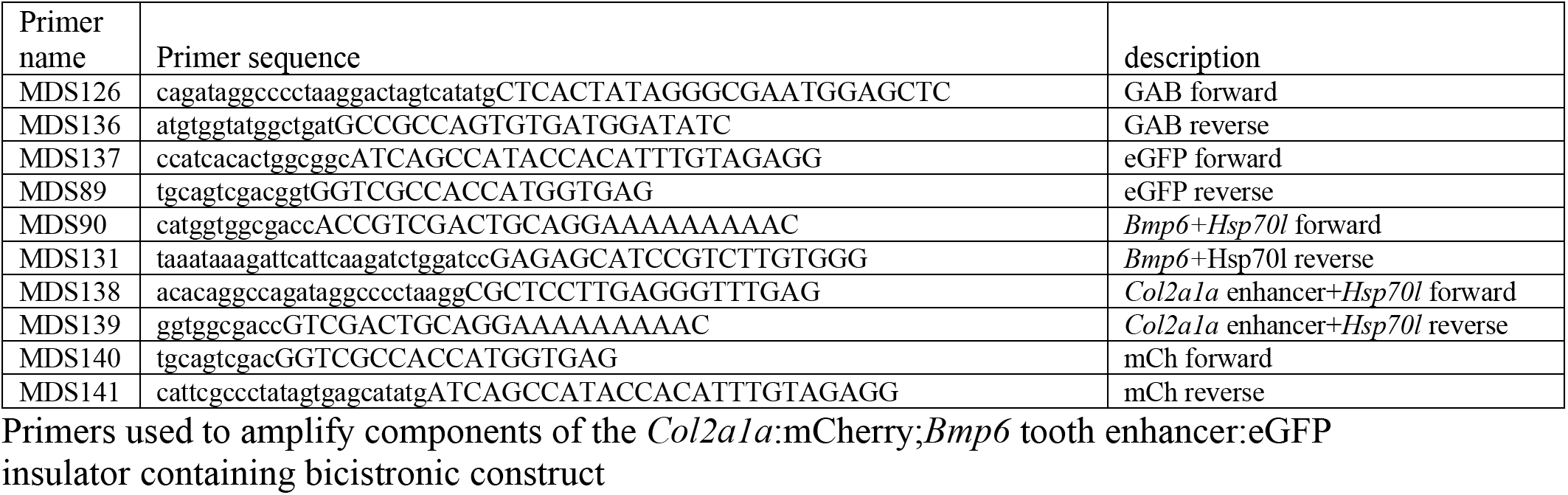

#### *Col2a1a* containing insulator construct #2

##### *Col2a1a* enhancer/*Hsp70l*àeGFP+GAB+mChß*Hsp70l*/*Bmp6* enhancer

The assembly of the second *Col2a1a* containing bicistronic construct is nearly identical to the first. All steps are the same except primers MDS137/89 were used to amplify mCherry in the first assembly step and primers MDS140/141 were used to amplify eGFP in the second assembly step. Due to identical sequence at the transition from *Hsp70l* to mCherry/eGFP and at the 3’ end of the SV40 polyA sequence for each reporter, the same primers can be used to amplify both off of the original reporter plasmids.

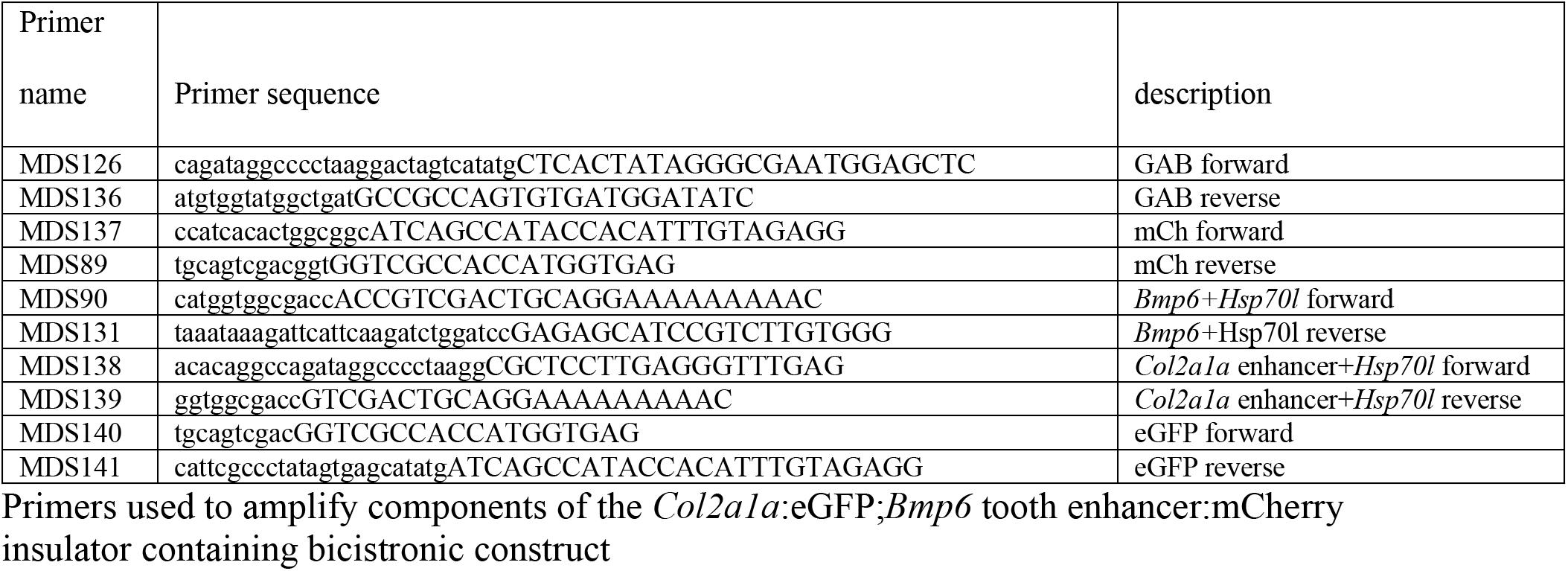

#### *Bmp6* intron 4 enhancer containing insulator construct

##### *Marine Bmp6* enhancer/*Hsp70l*àeGFP+GAB+mChß*Hsp70l*/Freshwater *Bmp6* enhancer

The first assembly step was the same as the previous two constructs, except the primer pair MDS90/131 was used to specifically amplify the freshwater *Bmp6* enhancer. Linearization of the plasmid and Gibson Assembly was completed as before. The resulting plasmid was digested with *Nde*I and *Bsu36*I and the fragments for the second half, Marine *Bmp6* enhancer/*Hsp70l* and mCherry, were amplified with MDS164/139 and MDS140/141 respectively. The newly digested plasmid and amplicons were combined with Gibson Assembly master mix and incubated following the manufacturer’s protocol.

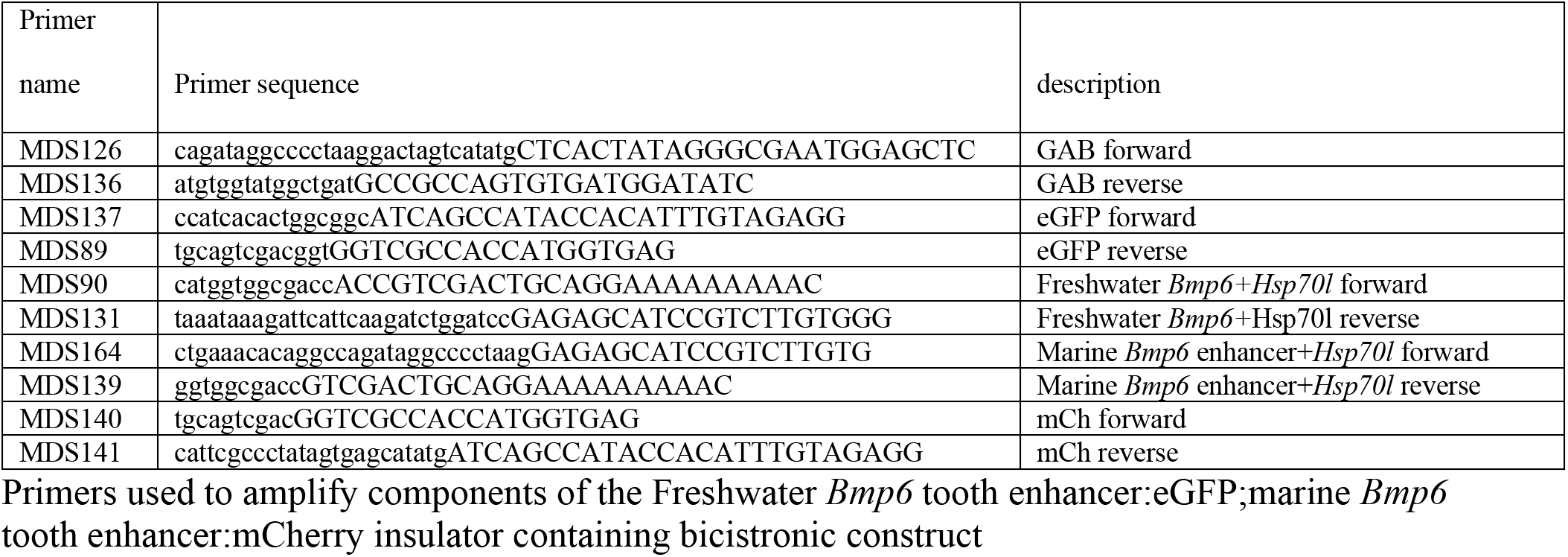

#### Scoring effectiveness of insulators

To assess insulator effectiveness, all surviving injected fish were raised to 7 days post fertilization. At this time point the *Bmp6* intronic enhancer drives robust reporter expression in multiple domains including the distal edges of the median and pectoral fins, while the *Col2a1a* enhancer drives expression in the notochord (Cleves et al., 2018; Erickson et al., 2016). Four anatomical domains were scored for insulator effectiveness: the left and right pectoral fins, the median fin, and the notochord. Insulator efficiency was scored on a scale of 0 (apparent complete lack of insulation) to 2 (fully insulated enhancers) for each domain in which expression was observed. Insulation activity was only assessed for domains in which expression of at least a single fluorophore was present. Since effectiveness was scored in F_0_ fish which are mosaic for the injected transgene, not all domains expressed a fluorophore.

### Supplemental Results

#### Insulator effectiveness in bicistronic constructs

Insulator scores were not significantly different across injection clutches for the *Col2a1a R2*:mCherry; *Bmp6* tooth enhancer:eGFP construct (Kruskal-Wallis left pectoral fin *P* = 0.075, right pectoral fin *P* = 0.52, median fin fold *P* = 0.116, Wilcoxon rank sum notochord *P* = 0.25), nor the *Col2a1a R2*:eGFP; *Bmp6* tooth enhancer:mCherry construct (Wilcoxon rank sum left pectoral fin *P* = 0.144, right pectoral fin *P* = 0.134, median fin fold *P* = 0.211), suggesting that the inter-clutch variation did not have a significant impact on insulation scores. The left pectoral fin (*P =* 0.036) and the median fin fold (*P* = 0.016) were found to be significantly different between the two constructs while the right pectoral fin (*P =* 0.68) and notochord (*P* = 0.29) were not significantly different.

#### Marine enhancer activity in the epithelium differs across tooth stage and fish size

In post-tooth number divergence fish activity of the freshwater enhancer was observed in the epithelium in both ventral and dorsal tooth plates in all pre-eruption teeth (marine:mCherry;freshwater:eGFP ventral: 59/59, dorsal: 44/44, and marine:eGFP;freshwater:mCherry ventral: 39/39, dorsal: 48/48), while the marine allele was observed in a subset of pre-eruption teeth (marine:mCherry;freshwater:eGFP ventral: 50/59 [84.7%], dorsal: 33/44 [75.0%], and marine:eGFP;freshwater:mCherry ventral: 35/39 [89.7%], dorsal: 34/48 [70.8%]). A higher percentage of early stage pre-eruption germs had marine activity in the epithelium compared to middle stage pre-eruption germs (marine:mCherry;freshwater:eGFP ventral: 20/23 [87.0%], dorsal: 24/29 [82.8%], and marine:eGFP;freshwater:mCherry ventral: 21/23 [91.3%], dorsal: 18/24 [75%]) than middle stage germs (marine:mCherry;freshwater:eGFP ventral: 30/36 [83.3%], dorsal: 9/15 [60.0%], and marine:eGFP;freshwater:mCherry ventral: 14/16 [87.5%], dorsal: 16/24 [66.7%]). In contrast to post-divergence, or > 20 mm total length, the marine enhancer in pre-divergence fish was active in every pre-eruption tooth germ (marine:mCherry;freshwater:eGFP ventral: 31/31, dorsal: 36/36, and marine:eGFP;freshwater:mCherry ventral: 29/29, dorsal: 29/29).

#### Mesenchymal bias differs across tooth stage, plate, and fish size

Mesenchymal bias, in which one enhancer was observed to drive a broader domain within the mesenchyme, was scored for post divergence fish. In early and middle stage teeth, we observed a consistent marine enhancer bias in the ventral (marine:mCherry;freshwater:eGFP early: 23/23, middle: 36/36, marine:eGFP;freshwater:mCherry early: 21/23 [91.3%], middle: 16/16) and dorsal tooth plates (early: 28/29, 96.6%, middle:15/15, marine:eGFP;freshwater:mCherry early: 24/24, middle: 24/24)). A larger proportion of functional, erupted teeth were observed to have a marine bias in the mesenchyme in the ventral tooth plate (marine:mCherry;freshwater:eGFP 87/108 [80.6%], marine:eGFP;freshwater:mCherry 86/97 [88.7%]) compared to the dorsal tooth plate (marine:mCherry;freshwater:eGFP 59/105 [56.2%], marine:eGFP;freshwater:mCherry 74/101 [73.3%] (Figure 5B-C). There was a reduction in the proportion of erupted teeth with a marine bias when comparing post to pre divergence fish for all integrations and tooth plates (pre-divergence marine:mCherry;freshwater:eGFP ventral: 54/80 [67.5%] and marine:eGFP;freshwater:mCherry ventral: 63/98 [64.3%], dorsal pre: 51/103 [49.5%]) (Figure 5B) except for the dorsal tooth plates in the freshwater:eGFP;marine:mCherry genotype (pre: 55/91 [60.4%], post: 59/105[56.2%]).

